# Rim aperture of autophagic membranes balances cargo inclusion with vesicle maturation

**DOI:** 10.1101/2023.06.01.543228

**Authors:** Oren Shatz, Milana Fraiberg, Alexandra Polyansky, Eyal Shimoni, Tali Dadosh, Sharon Wolf, Zvulun Elazar

**Author notes:** Corresponding author - Zvulun Elazar Department of Biomolecular Sciences The Weizmann Institute of Science 76100 Rehovot, Israel.

## Abstract

Autophagy eliminates cytoplasmic material by engulfment in membranous vesicles targeted for lysosome degradation. Nonselective autophagy coordinates the sequestration of bulk cargo with the growth of the isolation membrane (IM) in a yet-unknown manner. Here we show the yeast IM expands while maintaining a rim sufficiently wide for sequestration of large cargo but tight enough to mature in due time. An obligate complex of Atg24/Snx4 with Atg20 or Snx41 assembles locally at the rim in a spatially-extended manner that specifically depends on autophagic PI(3)P. This assembly stabilizes the open rim to promote autophagic sequestration of large cargo in correlation with vesicle inflation. Moreover, constriction of the rim by the PI(3)P-dependent Atg2-Atg18 complex and clearance of PI(3)P by Ymr1 antagonize the rim opening to promote autophagic maturation and consumption of small cargo. Tight regulation of membrane rim aperture by PI(3)P thus couples the mechanism and physiology of nonselective autophagy.

## Introduction

Autophagy is a conserved catabolic process that targets cytoplasmic portions to lysosomal degradation by sequestration in an isolation membrane (IM). Upon autophagic signaling, the IM forms *de novo*, expands, and seals into a double-membrane vesicle termed autophagosomes. Maturation of autophagosomes by clearance of associated autophagic proteins is then followed by fusion with the lysosome and degradation of engulfed cargo (Dikic and Elazar, 2018). The selective autophagosome exclusively engulfs specific cargo to be eliminated in a process that tightly couples IM expansion to close interaction of IM-conjugated proteins of the ATG8 superfamily with cargo-bound receptor proteins (Zaffagnini and Martens, 2016). In contrast, nutrient starvation leads to formation of autophagosomes loaded with apparent bulk cytoplasmic content (Baba et al., 1994). Genetic screens coupled with *in vitro* studies have identified and characterized most proteins and lipids participating in nonselective autophagosome biogenesis (Hu and Reggiori, 2022). Still, the morphological features and coordination of the process with cargo sequestration remain unclear.

Molecular dissection of the differential contribution of autophagy proteins to intermediate steps in autophagosome biogenesis has been experimentally challenging, as complete loss of core autophagy protein activity leads to the arrest of the process at an early stage. Indeed, even downstream-acting factors like the Atg2-Atg18 complex (Suzuki et al., 2007) are nevertheless essential for IM expansion (Noda, 2021). Alternative approaches for quantitative down-regulation of protein function have so far indicated quantitative roles for Atg9 and the Atg8 system in membrane formation and determination of autophagosome size (Jin and Klionsky, 2014), respectively. Quantitative contribution to intermediate steps, however, remains speculative. Interestingly, recent mutational analysis of Atg2 revealed a later role in the proper positioning of Atg9 at the bridge between IM rim and associated endoplasmic reticulum, characterized in a selective cargo model (Gomez-Sanchez et al., 2018).

Similarly challenging is the morphological dissection of IM expansion, given the diffraction-limited resolution of the IM and associated factors under live fluorescence microscopy (Geng et al., 2008), whereas in the genetically-tractable budding yeast model system electron microscopy often captures the terminal state of IM expansion (Baba *et al*., 1994). Alternatively, the selective giant Ape1 complex (GAC) cargo has been employed for static spatially-informed visualization of IM and associated factors (Suzuki et al., 2013) and more recently in kinetic characterization of IM expansion (Schutter et al., 2020). Still, the implications for sequestration of bulk cargo remain unaddressed. In addition, the mechanistic relevance of GAC-informed spatial patterns is uncertain, taking into account the modulation by selectivity scaffold Atg11 of the nonselective process (Sekito et al., 2009) – for which it is dispensable (Shintani and Klionsky, 2004).

Here we establish an experimental system to combine volumetric visualization of Atg11-independent, strictly nonselective autophagosome biogenesis under fluorescence and correlated electron microscopy with quantitative and timely down-regulation of late-acting autophagy proteins. We demonstrate that the nonselective IM expands while maintaining a rim sufficiently wide to allow the sequestration of large cargo but tight enough to mature in due time. PI(3)P-dependent local assembly of a spatially-extended, autophagy-specific obligate complex of Atg24/Snx4 with Atg20 or Snx41 stabilizes the rim to promote vesicle inflation and sequestration of large cargo. The reciprocal rim constriction activity of the PI(3)P-dependent Atg2-Atg18 complex and PI(3)P clearance activity of Ymr1 promote maturation and flux of small cargo. Our observations thus suggest that tuning these opposing activities balances vesicle maturation with cargo size. Regulation of rim aperture through PI(3)P effectors thus couples the mechanism and physiology of nonselective autophagy.

## Results

### Nonselective isolation membranes (IMs) expand as large vesicles with a narrow rim

The power of genetics in budding yeast has been instrumental in identification and visualization of proteins that associate with the selective (Suzuki *et al*., 2013) and nonselective (Graef et al., 2013) isolation membrane (IM). Still, the morphological features of nonselective autophagosome biogenesis and their physiological relevance have been largely overlooked. Here we characterized the strictly-nonselective autophagy by elimination of the selectivity scaffold Atg11 (Zientara-Rytter and Subramani, 2020). To this end, we visualized the IM by Atg8 (Kirisako et al., 2000), PI(3)P – a mechanistically-important phospholipid previously localized to autophagic membranes (Cheng et al., 2014) – by the phox-homology (PX) domain of Vam7 (PX^Vam7^) (Cheever et al., 2001), vacuolar membrane (VM) by Pho8 (Klionsky and Emr, 1989), and the endoplasmic reticulum (ER) by Sec61 (Wilkinson et al., 1996). Upon induction of autophagy by the mTOR inhibitor rapamycin (Noda and Ohsumi, 1998) in wildtype cells, spherical Atg8-positive structures appeared between the VM and ER, where PI(3)P marker PX^Vam7^ was visibly enriched (Figure 1A) – suggesting these are isolation membranes. As a negative control, knock-out of the phagophore-ER tether and lipid transporter Atg2 (Gomez-Sanchez *et al*., 2018; Kotani et al., 2018; Osawa et al., 2019) arrested expansion and enrichment with PI(3)P of these structures (Figure 1A). Moreover, these structures also exhibited the reported colocalization pattern of autophagic marker proteins Atg1, Atg2, and Atg5 on nonselective phagophores throughout the IM surface, at the IM base and IM rim, respectively (Figure S1A) (Graef *et al*., 2013). We therefore conclude that these are indeed *bona fide* isolation membranes and that, in our hands, nonselective IMs assume a spherical shape.

**Figure 1:**
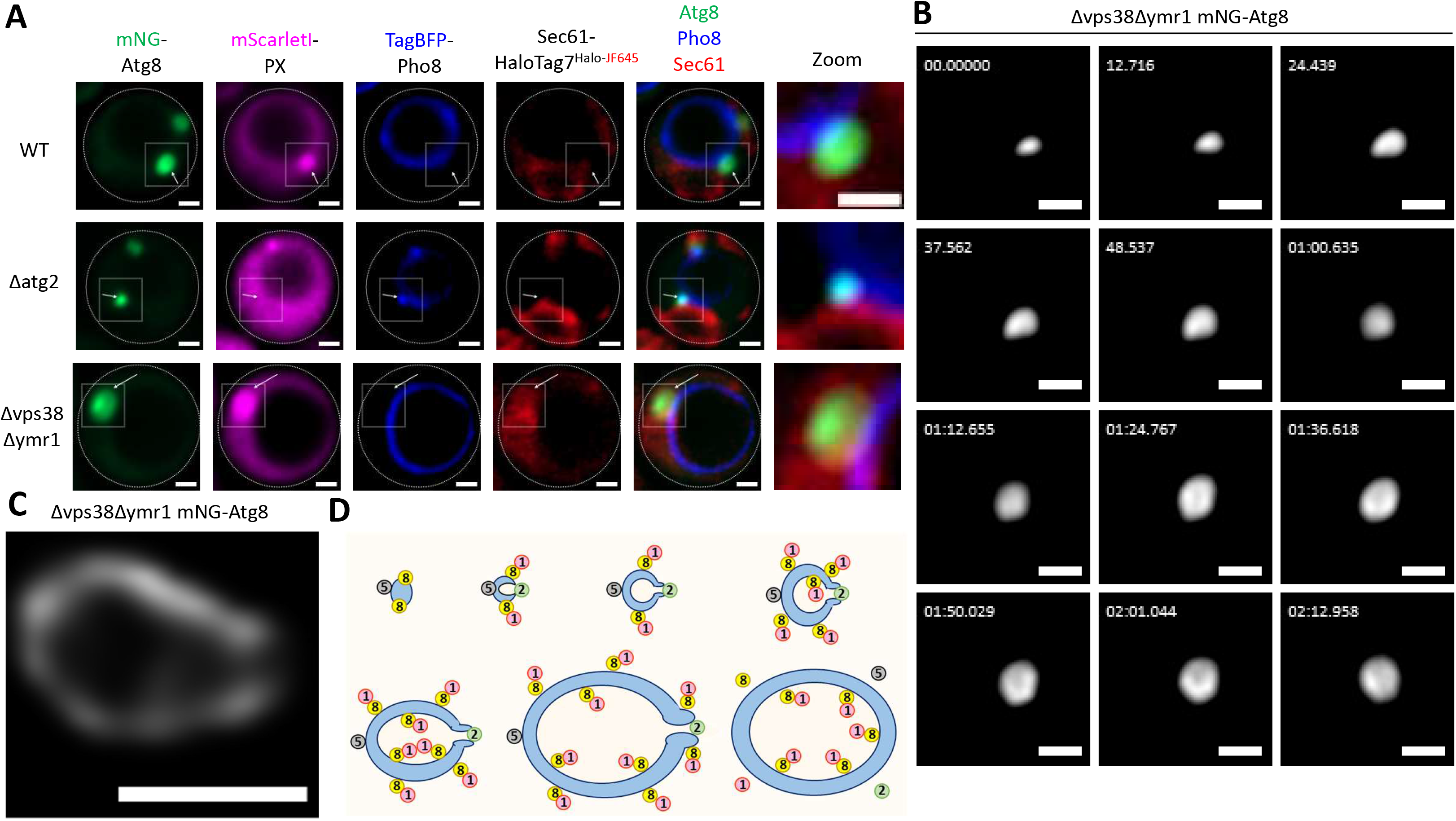
Nonselective isolation membranes (IMs) expand as large vesicles with a narrow rim. (A) Spherical Atg8/PI(3)P double-positive isolation membranes (IMs) appear between the vacuolar membrane and endoplasmic reticulum upon induction of nonselective autophagy by rapamycin in absence of Atg11. Cells of indicated genotype expressing mNG-Atg8, TagBFP-Pho8, Sec61-HaloTag7, and mScarletI-PX^Vam7^ were grown to logarithmic phase in presence of 10μM CuSO_4_, stained with Halo-JF645, treated for 2.5-3h with rapamycin and PMSF and imaged by widefield microscopy. Scale bar 1μm. (B) Timelapse microscopy of IM expansion in PI(3)P-augmented conditions shows an increase in vesicle size with the maintenance of a narrow rim. Δvps38Δymr1 cells expressing mNG-Atg8 were grown to logarithmic phase, treated for 1.5-3h with rapamycin, and imaged by widefield microscopy at indicated time-points. Scale bar 1μm. (C) Visually-enhanced PI(3)P-augmented IM exhibits a large body and a narrow rim. Δvps38Δymr1 cells expressing mNG-Atg8 were grown to logarithmic phase, treated for 4h with rapamycin, and imaged by Leica confocal microscopy with Lightning processing. Scale bar 1μm. (D) Schematic representation – nonselective isolation membranes first bend from a flat disc into a small cup, whose rim is maintained by Atg2. Then they expand into spherical membrane as a vesicle that increases in size while maintaining a narrow rim. Membrane-associated autophagy proteins are indicated by numbers: Atg1, Atg2, and Atg5 localize throughout the IM surface, at the IM base, and at its rim, respectively. Maturation follows by sealing of the single continuous isolation membrane into an autophagosomal double membrane and clearance of proteins associated with the outer membrane, while Atg1 remains associated with Atg8 covalently bound to the inner membrane.

Their strictly-closed appearance of IMs led us to explore the possibility that nonselective IMs maintain a narrow rim even during vesicle expansion. PI(3)P supports both the membrane conjugation of Atg8 through the PI(3)P effector Atg21 (Juris et al., 2015) and the recruitment of Atg2 through interaction with the PI(3)P effector Atg18 (Kobayashi et al., 2012; Obara et al., 2008). We sought to augment rim aperture and the vesicle size by supporting higher autophagy-specific PI(3)P levels. We therefore knocked out the PI(3)P phosphatase Ymr1, previously implicated in autophagy (Cebollero et al., 2012), and the endosomal PI3-kinase complex II specificity subunit Vps38 (Kihara et al., 2001). Induction of autophagy in Δvps38Δymr1 cells led to formation of large IMs with narrow rim (Figure 1A) – better resolved under Leica HyVolution confocal microscopy (Figure 1C). As for wildtype, Δvps38Δymr1 IMs were enriched with a PI(3)P marker (Figure 1A) and exhibited reported colocalization patterns of Atg1 and Atg5 (Figure S1A). Finally, time-lapse microscopy of both wildtype (Figure S1B) and Δvps38Δymr1 IMs (Figure 1B) demonstrated growth in vesicle size but not in rim aperture (Figure 1D). This is further supported by the observation of a spatially-constricted pattern of rim marker Atg2 in the course of autophagosome biogenesis (Figure S1C).

### The Atg2-Atg18 complex promotes constriction of the IM rim

Whereas cup-shaped phagophores are often ascribed a wide rim of unknown regulation, we predicted that constriction of the narrow rim observed in our hands might require tight regulation to coordinate sealing with sequestration of cargo. Considering the localization of Atg2 to the rim, we assayed the effect of partial Atg2 depletion on rim morphology by employing the auxin inducible degron (AID) approach (Morawska and Ulrich, 2013; Nishimura et al., 2009). As expected, titration of the auxin hormone indole-3-acetic acid (IAA) quantitatively depleted Atg2 tagged with a minimal degron (AID*) – in correlation with loss of autophagic flux, indicated by GFP-Atg8 cleavage (Figure S2A). As controls, wildtype cells and Atg2-AID*(P88L) cells carrying the auxin-desensitizing P88L mutation of the degron (Rouse et al., 1998) maintained autophagic flux, excluding non-specific effects of IAA on autophagy (Figure S2A). Treatment of wildtype cells with a non-saturating IAA concentration led to formation of round IMs in wildtype cells, whereas the partial depletion of Atg9, or knock-out of Atg2, Atg18 or Atg9 arrested membrane expansion or formation (Figure 2A) – as expected given their core roles in autophagy. Surprisingly, IMs with visibly-open rims were readily observed upon partial depletion of Atg2 and Atg18 (Figure 2A) from the IM rim (Figure S2B), supporting an *in situ* role in rim constriction. As above, the *bona fide* autophagic identity of these membranes is confirmed by their enrichment with PI(3)P marker PX^Vam7^ (Figure 2A), genetic requirement for core autophagy factors (Figure S2C) and expected localization of Atg1 and Atg5 (Figure S2D). Finally, correlative light electron microscopy (CLEM) analysis demonstrated a wide-open double-membrane in spatial correlation with fluorescent detection of Atg8 (Figure 2B, S2E, Movie M1).

**Figure 2:**
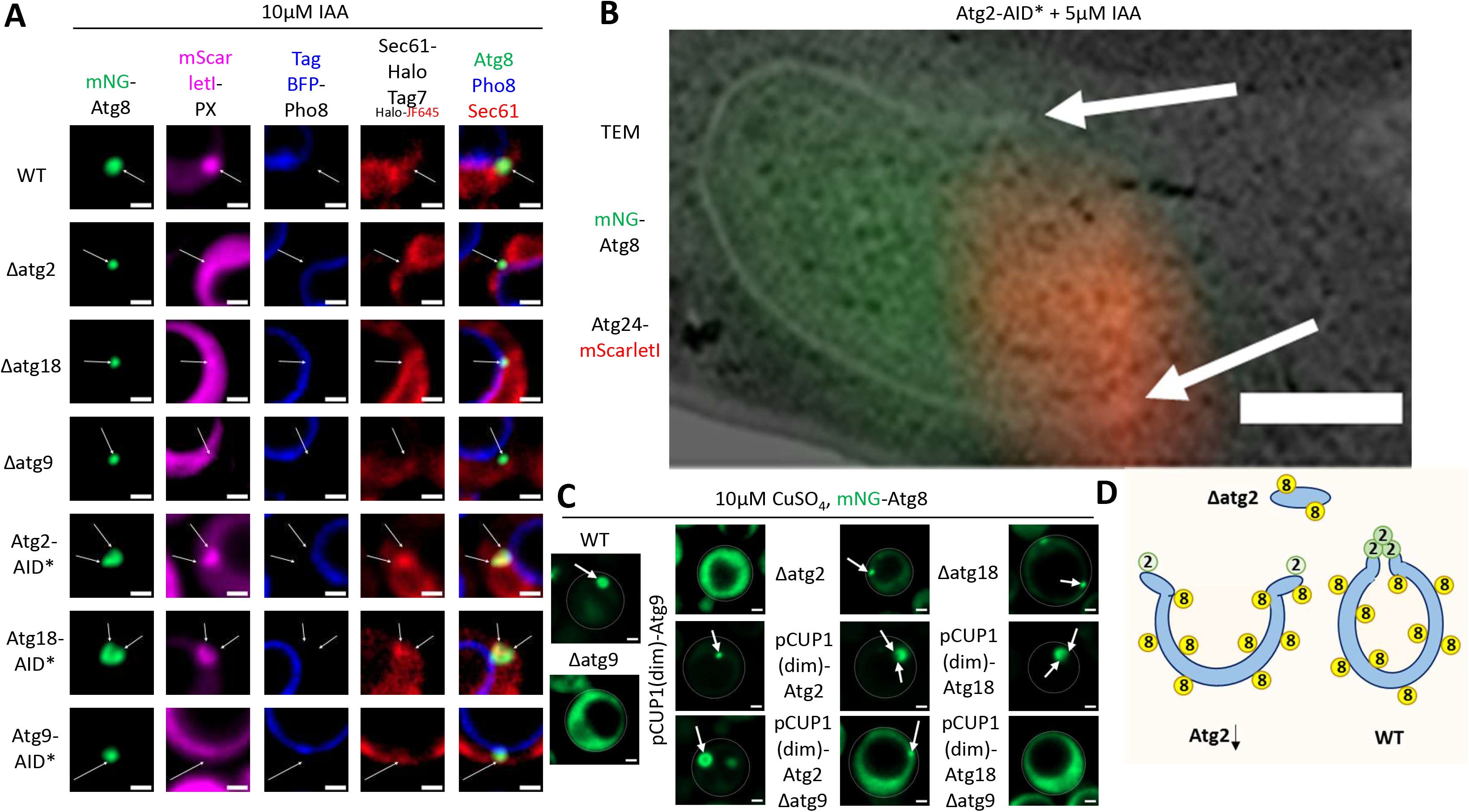
The Atg2-Atg18 complex promotes constriction of the IM rim. (A) Partial AID*-mediated depletion of Atg2 or Atg18 – but not Atg9 – leads to formation of IMs with wide-open rims. Cells of indicated genotype expressing mNG-Atg8, TagBFP-Pho8, Sec61-HaloTag7, and mScarletI-PX^Vam7^ were grown to logarithmic phase in presence of 10μM CuSO_4_, stained with Halo-JF645, treated for 2.5-3h with rapamycin, PMSF and 10μM IAA, and imaged by widefield microscopy. Scale bar 1μm. (B) Correlative light electron microscopy (CLEM) analysis of partially-depleted Atg2 IMs shows an open, locally-deformed Atg24-associated rim (arrows). Atg2-AID* cells expressing mNG-Atg8 and Atg24-mScarletI were grown to logarithmic phase and treated for 100min with rapamycin, PMSF, and 5μM before CLEM processing and imaging. A single z-slice of reconstructed tomography z-stack is shown. Scale bar 200nm. (C) Partial transcriptional down-regulation of Atg2 or Atg18 – but not Atg9 – using a weak version of the copper-induced promoter CUP1 (pCUP1(dim)) leads to the formation of IMs with open rims. Cells of indicated genotype expressing mNG-Atg8 were grown to logarithmic phase in presence of 10μM CuSO_4_, treated for 2.5h with rapamycin and PMSF, and imaged by widefield microscopy. Scale bar 1μm. (D) Schematic representation – wildtype levels of local Atg2 activity facilitate rim constriction at the course of IM expansion, while partially active Atg2 maintains membrane expansion but compromises rim constriction.

As independent means of protein down-regulation, we replaced the native promoters of Atg2 or Atg18 with pCUP1(dim) – a weak variant of the copper-inducible CUP1 promoter (Rugbjerg et al., 2015). Western blot analysis confirms that in absence of copper, this promoter dramatically reduces the expression of Myc-tagged Atg2 in correlation with loss of flux – both restored upon induction by copper (Figure S2F). Open IMs were observed upon mild copper treatment of pCUP1(dim)-promoted Atg2 or Atg18 – but not in wildtype cells, where circular membranes were observed, or upon complete loss of Atg2, Atg18 or Atg9 – where IMs failed to form or develop (Figure 2C). Importantly, partial down-regulation of Atg9 variably led to an arrest in membrane formation or expansion – or to formation of wildtype-shaped round IMs (Figure 2C). This result is in line with the reported quantitative role of Atg9 in vesicle nucleation (Jin et al., 2014). While Atg9 interacts with Atg2 locally at the rim (Gomez-Sanchez *et al*., 2018), our observation, however, excludes a quantitative morphogenic role for Atg9 therein. Our observations thus indicate that the Atg2-Atg18 complex promotes constriction of the IM rim, whereas inadequate activity of rim-localized Atg2 leads to expansion of aberrantly-open IMs (Figure 2D).

### An obligate complex of Atg24 with Atg20 or Snx41 assembles at the IM rim

The role of Atg2 in rim constriction in particular and IM shaping, in general, led us to consider the regulatory activity of other factors at the IM rim. The endosomal sorting nexin (SNX)-Bin/Amphiphysin/Rvs (BAR) protein Snx4/Atg24 and its distinct SNX-BAR interactors Atg20/Snx42 and Snx41 (Ma et al., 2017) are recruited through their PX domain to PI(3)P-positive membranes (Hettema et al., 2003) and associate through their BAR domains with curved membranes (Peter et al., 2004). Moreover, the Atg24-Atg20 complex is implicated in autophagy by association with the selective Cvt structure (Nice et al., 2002). In light of the outward-facing local membrane deformation at the rim observed above upon partial depletion of Atg2 (Figure 2B) and recently reported in wildtype conditions (Bieber et al., 2022), we asked whether an Atg24 complex with Atg20 or Snx41 may play an autophagy-specific role at the IM rim.

Upon induction of autophagy Atg24 localized to the IM rim, but mislocalized upon loss of its BAR domain-containing C-terminal half (amino acids 172-423), its PX domain-containing N-terminal half (amino acids 2-171) or the autophagy specificity PI3-kinase subunit Atg14 (Figure 3A, 3B) – in line with autophagic recruitment of Atg24 by both autophagic PI(3)P and local membrane curvature. Importantly, Atg24 also mislocalized from the rim when Atg20-AID* was transiently depleted in absence of Snx41, or when Snx41-AID* was depleted in absence of Atg20. Nevertheless, Atg24 maintained its localization when only one of its co-factors was missing (Figure 3B) – suggesting the formation of a local obligate Atg24 complex (termed Atg24C hereafter) with Atg20, Snx41 or both. Accordingly, Atg20 localized to the rim in wildtype or Snx41-deficient cells but mislocalized upon transient or constitutive loss of Atg24 (Figure 3C, S3A), while the rim localization of Snx41 reciprocally depended on Atg24 but not Atg20 (Figure S3B, S3C).

**Figure 3:**
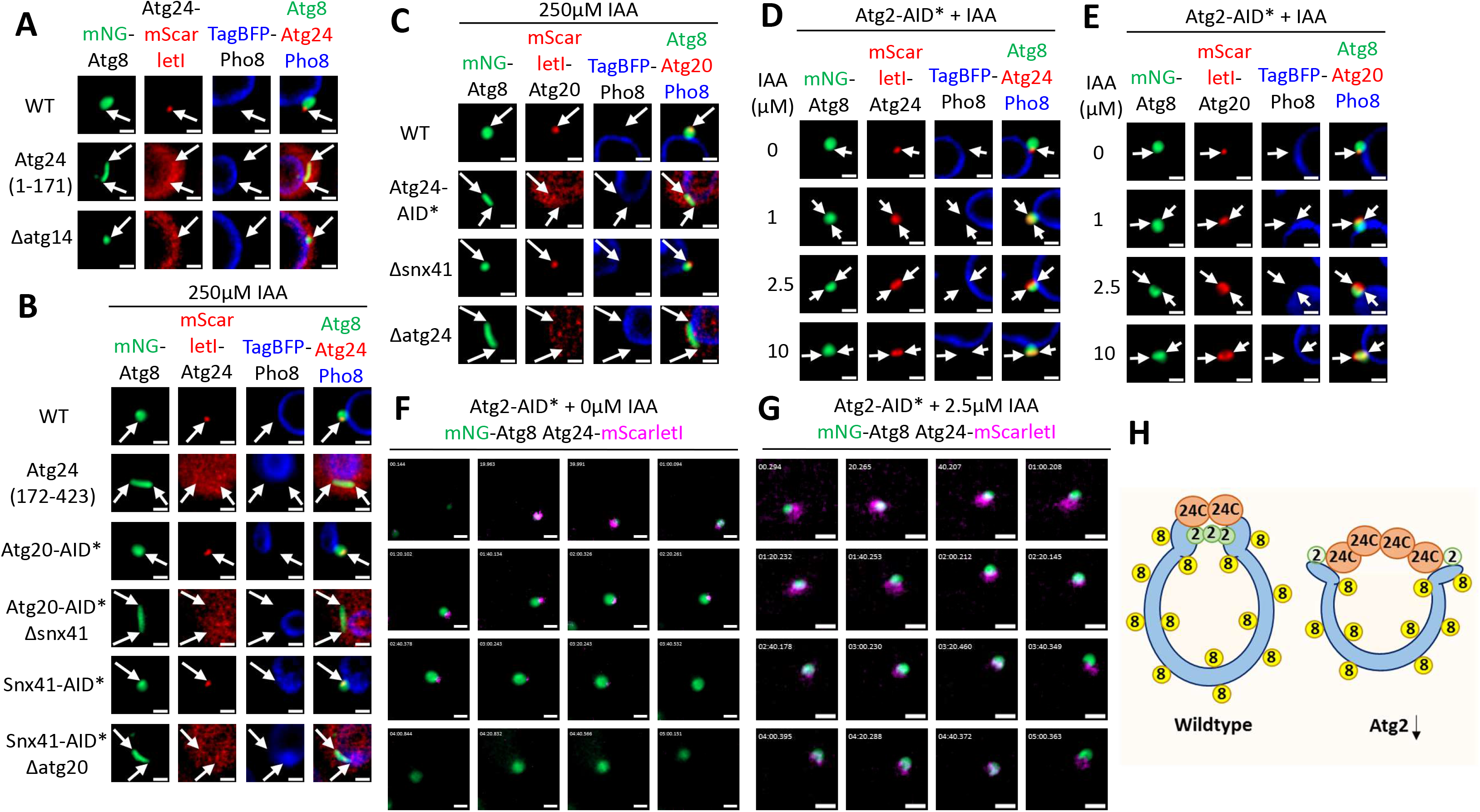
An obligate complex of Atg24 with Atg20 or Snx41 assembles at the IM rim. (A) Atg24 C-terminally tagged with mScarletI localizes to the IM rim in a manner dependent on its N-terminal PX domain and autophagic PI(3)P. Cells of indicated genotype expressing mNG-Atg8, TagBFP-Pho8, and Atg24-mScarletI were grown to logarithmic phase, treated for 2.5-3.5h with rapamycin and PMSF, and imaged by widefield microscopy. Scale bar 1μm. (B) Atg24 N-terminally tagged with mScarletI localizes to the IM rim in a manner dependent on its C-terminal BAR domain and either Atg20 or Snx41. Cells of indicated genotype expressing mNG-Atg8, TagBFP-Pho8, and mScarletI-Atg24 were grown to logarithmic phase, treated for 2-4h with rapamycin and PMSF, as well as 250μM IAA for the depletion of indicated AID*-tagged proteins, and imaged by widefield microscopy. Scale bar 1μm. (C) Atg20 N-terminally tagged with mScarletI localizes to the IM rim in a manner dependent on Atg24 but not Snx41. Cells of indicated genotype expressing mNG-Atg8, TagBFP-Pho8, and mScarletI-Atg20 were grown to logarithmic phase, treated for 2.5-4h with rapamycin and PMSF, as well as 250μM IAA for depletion of indicated AID*-tagged proteins, and imaged by widefield microscopy. Scale bar 1μm. (D, E) Atg24 (D) and Atg20 (E) increase their prominence at the IM rim in correlation with AID*-mediated loss of Atg2. Atg2-AID* cells expressing mNG-Atg8, TagBFP-Pho8 and mScarletI-tagged Atg20 or Atg24 were grown to logarithmic phase and treated for 2-3h with rapamycin, PMSF and indicated IAA concentration, before imaging by widefield microscopy. Scale bar 1μm. (F, G) Atg24 is associated with the narrow rim of non-depleted (F) and open rim of partially depleted (G) Atg2-AID* IMs. Atg2-AID* cells expressing mNG-Atg8 and Atg24-mScarletI were grown to logarithmic phase, treated for 2-3h with rapamycin and 0μM (F) or 2.5μM (G) IAA, and imaged by widefield microscopy at indicated time-points. Scale bar 1μm. (H) Schematic representation – an obligate complex of Atg24 (Atg24C, or 24C in short) with Atg20 or Snx41 localizes in a constricted manner to the narrow rim of the wildtype IM or in a spatially-extended manner to the open rim of partially-depleted Atg2-AID* IM.

As partial loss of Atg2-Atg18 activity was shown above to open the IM rim (Figure 2), the formation of an Atg24 complex would be expected to decorate the wider PI(3)P-positive curved surface exposed under such conditions. Indeed, partial depletion of Atg2-AID* led to spatially-extended decoration of the rim surface by Atg24 (Figure 2B, S3D) and Atg20 (Figure S3E) in an Atg18 and Atg14-dependent manner, respectively. Moreover, quantitative titration of Atg2-AID* leads to a gradual increase in prominence of rim-localized Atg24 (Figure 3D) and Atg20 (Figure 3E) – quantitatively linking rim dysregulation due to loss of Atg2-Atg18 activity to a spatial extension of the Atg24-Atg20 complex assembly. Finally, timelapse imaging shows that upon basal expression of Atg2 (tagged with AID* but mock-treated), Atg24 initially surrounds the edges of the unexpanded IM disc but then rapidly redistributes to the nascent narrow IM rim in a spatially-constricted and temporally-restricted manner before clearance at maturation (Figure 3F). Partial depletion of Atg2-AID*, however, leads to spatially-extended localization of Atg24 throughout the wide-open rim of the growing membrane (Figure 3G) – where it remains stably associated (Figure S3F). In conclusion, Atg24 forms an autophagy-specific spatially-extended obligate complex with Atg20 (or Snx41) at the IM rim throughout the course of membrane expansion until maturation, based on interactions with PI(3)P and with local membrane curvature (Figure 3H). This is in line with the recently-reported cooperative binding and remodeling activity of the Atg24-Atg20 heterodimer – but not the Atg24-Atg24 homodimer – towards autophagosome-like membranes *in vitro* (Reinhart et al., 2022).

### The Atg24 complex promotes spherical IM morphogenesis

Intriguingly, we observed a consistent morphological change of the isolation membrane from circular to elongated upon loss or mislocalization of Atg24 (Figure 3A-C, S3A-C). Accordingly, in our hands, transient depletion of Atg24-AID* or constitutive loss of Atg24 led to the formation of elongated Atg8-positive membranes (Figure 4A). Their *bona fide* autophagic identity is attested by their enrichment with PI(3)P marker PX^Vam7^ (Figure 4A), the canonical localization of Atg1, Atg2 and Atg5 throughout their surface, at their rim and at their base, respectively (Figure 4B) and their loss upon further knock-out of core autophagy genes (Figure S4A). Importantly, similar morphology is observed in Atg11-deficient cells of the SEY6210 and BY4741 background – excluding a w303 background-specific effect of Atg24C (Figure S4B). Leica HyVolution STED analysis of Δvps38Δymr1Δatg24 cells suggests these elongated IMs are not flat single-membrane ribbons but rather narrow double-membrane tubes (Figure 4C). This is supported by the CLEM observation that wide-open IMs observed upon partial depletion of Atg2 in presence of Atg24 (Figure 2B, Movie 1) are reduced to tubular IMs of no visible rim in absence of Atg24 (Figure 4D, S4C, Movie 2). Finally, quantitative AID*-mediated depletion of Atg24 in Δvps38Δymr1 cells shows a graded morphogenic transition from spherical to the tubular (Figure S4D) and timelapse microscopy of Δvps38Δymr1Δatg24 cells shows gradually-elongating tubules (Figure 4E). Our data therefore implicate the rim-localized Atg24 complex in a spherical morphogenesis of the isolation membrane (Figure 4F).

**Figure 4:**
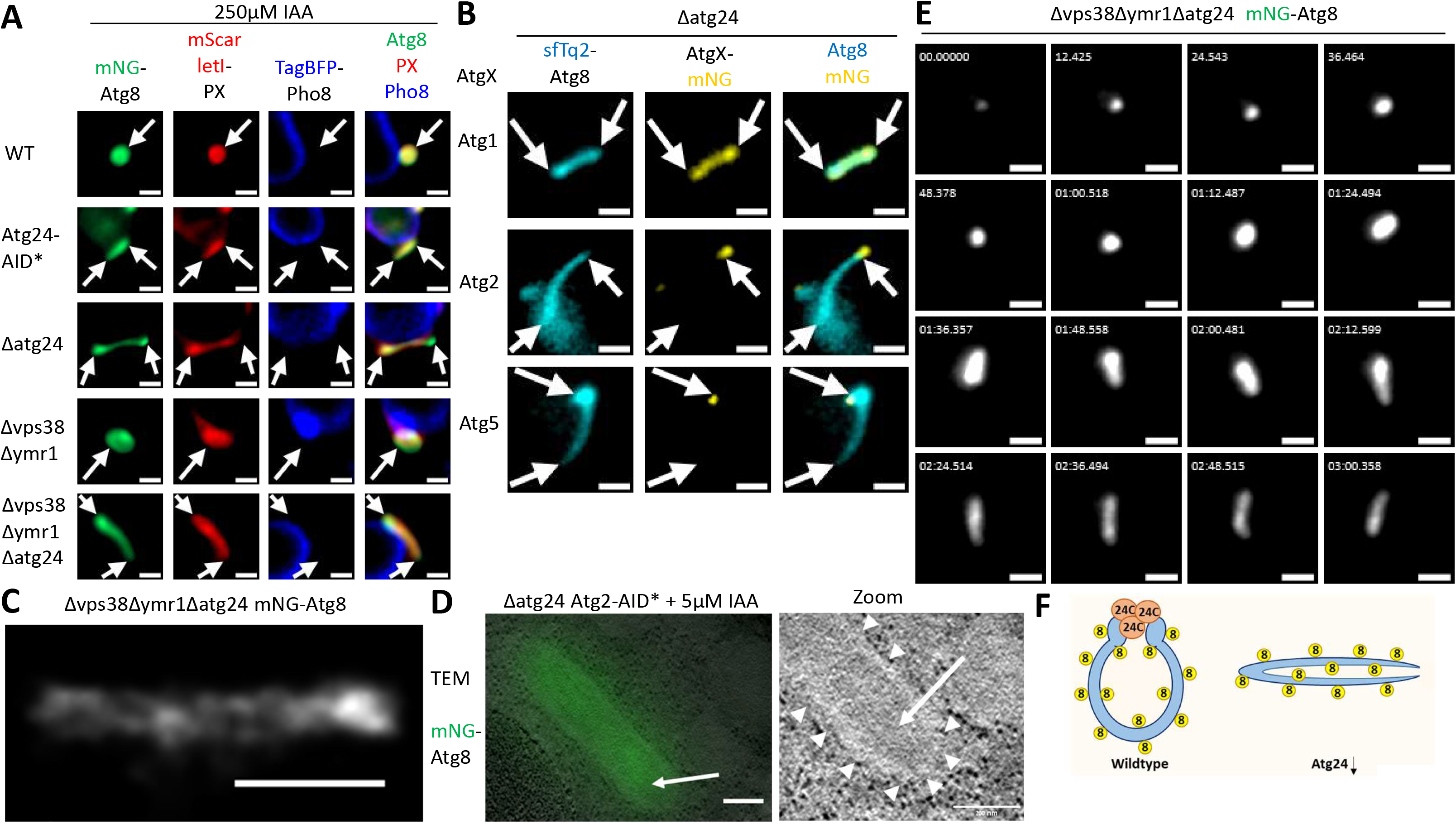
The Atg24 complex promotes spherical IM morphogenesis. (A) Loss of Atg24 leads to formation of elongated Atg8/PI(3)P double-positive IMs. Cells of indicated genotype expressing mNG-Atg8, mScarletI-PX^Vam7^, and TagBFP-Pho8 were grown to logarithmic phase in presence of 10μM CuSO_4_, treated for 1.5-2h with rapamycin, PMSF and 250μM IAA and imaged by widefield microscopy. Scale bar 1μm. (B) Canonical association pattern of autophagy proteins with the Atg24-deficient IM – Atg1, Atg2, and Atg5 localize throughout the IM surface, at its base, and at its rim, respectively. Wildtype and Δatg24 cells expressing sfTq2-Atg8 and indicated mNG-tagged autophagy factors were grown to logarithmic phase, treated for 1-2h with rapamycin and PMSF, and imaged by widefield microscopy. Scale bar 1μm. (C) Leica HyVolution STED microscopy identifies Atg24-deficient IMs as narrow tubes. Δvps38Δymr1Δatg24 cells expressing mNG-Atg8 were grown to logarithmic phase, treated for 2.5h with rapamycin, and imaged by Leica HyVolution STED microscopy. Scale bar 1μm. (D) CLEM analysis of Atg24-deficient cells exhibits narrow tubular Atg8-positive IMs (arrowheads) whose interior (arrow) excludes cytosolic particles like ribosomes. Δatg24 Atg2-AID* cells expressing mNG-Atg8 were grown to logarithmic phase and treated for 100min with rapamycin, PMSF, and 5μM before CLEM processing and imaging. A single z-slice of reconstructed tomography z-stack is shown. Scale bar 200nm. (E) Timelapse microscopy of IM expansion in PI(3)P-augmented, Atg24-deficient conditions show gradual elongation of a narrow IM. Δvps38Δymr1Δatg24 cells expressing mNG-Atg8 were grown to logarithmic phase, treated for 3h with rapamycin, and imaged by widefield microscopy at indicated time-points. Scale bar 1μm. (F) Schematic representation – Atg24C supports a spherical shape during IM expansion, whereas a tubular IM forms in its absence.

### Spatially-extended assembly of the Atg24 complex opens the IM rim

Our observations so far indicate that the Atg2-Atg18 complex and the Atg24 complex both localize to the IM rim and exert morphogenic effects: Atg2 promotes rim constriction (Figure 2) while Atg24C localizes to the rim (Figure 3) to promote spherical morphogenesis (Figure 4). As widening of the rim upon partial loss of Atg2 activity is correlated with spatial extension of the Atg24 complex on the rim surface (Figure 2B, 3B, 3C), we decided to test whether the rim is actively opened by the complex. For this we analyzed the differential effects of Atg24-Atg20 distribution patterns on rim aperture upon partial depletion of Atg2. Mild AID*-mediated depletion of Atg2 led to a mildly-extended assembly of Atg24-mScarletI next to a mildly-open rim (Figure 5A). In comparison, both Atg24 assembly and rim aperture were visibly augmented upon fusion of Atg24 to superfolder Cherry 3V (sfCh3V), a full-length version of a recently-improved split fluorescent mCherry (Feng et al., 2019), or upon tethering of Atg24-mScarletI to the ER by the transmembrane domain of Sec71 (TMD^Sec71^) (Figure 5A), and a similar effect of sfCh3V was observed for Atg20 (Figure 5B). Importantly, Atg24 variants maintained their rim localization upon loss of Vps38 but mislocalized upon loss of Atg14 or combined loss of Atg20 and Snx41 (Figure S5A). These observations together indicate that artificial gain of spatially-extended distribution of autophagic Atg24C may promote excessive widening of the rim upon partial depletion of Atg2, ascribing an active rather than passive role for Atg24C at the open rim.

**Figure 5:**
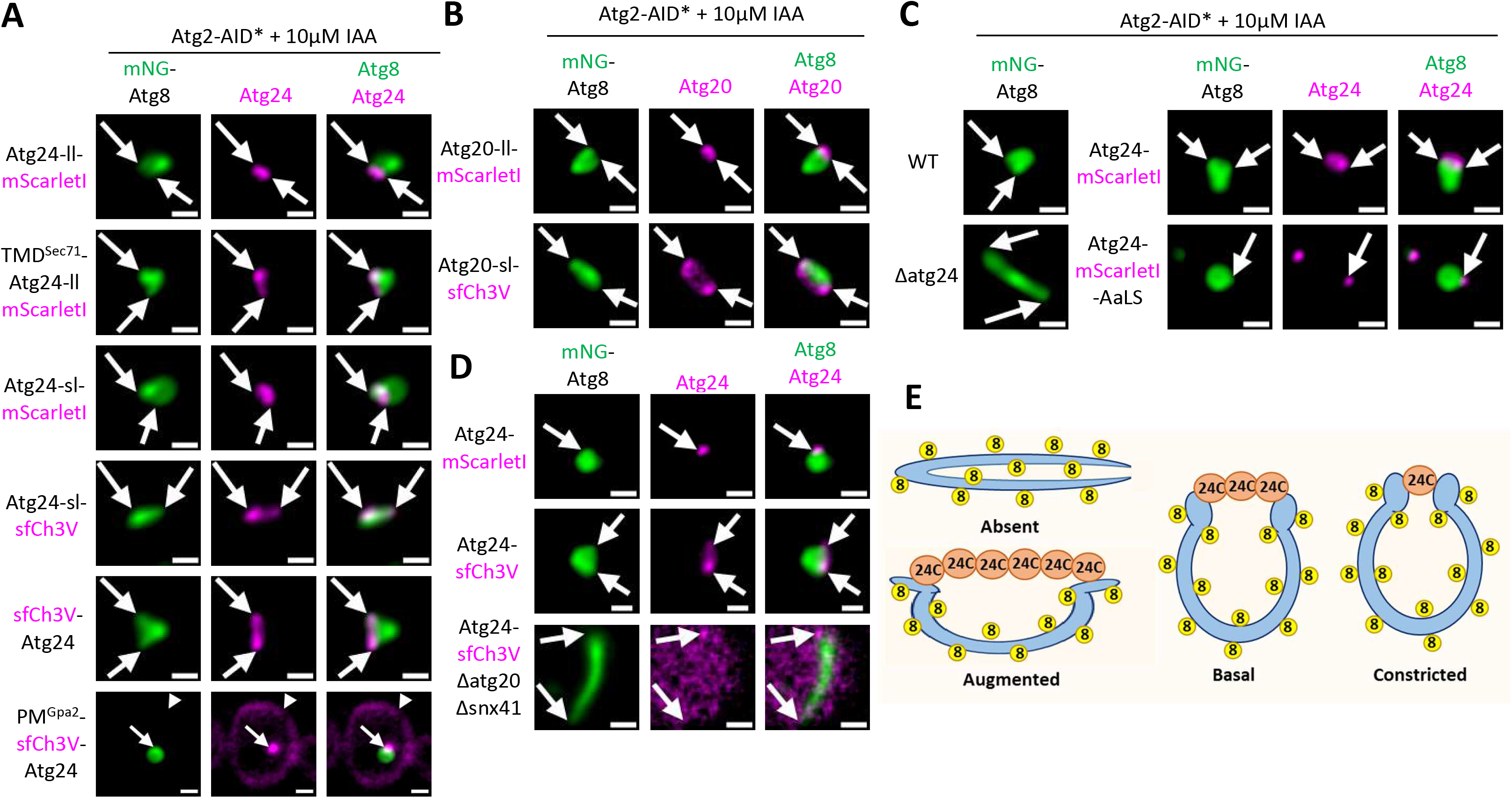
Spatially-extended assembly of the Atg24 complex opens the IM rim. (A, B) The spatial extension of Atg24 or Atg20 assemblies correlates with opening of the IM rim upon partial depletion of Atg2-AID*. Atg2-AID* cells of indicated genotype expressing mNG-Atg8 and indicated Atg24 (A) or Atg20 (B) alleles were grown to logarithmic phase in presence of 10μM CuSO4 and treated with rapamycin, PMSF, and 10μM IAA for 2.5-4h, before imaging by widefield microscopy. Arrowhead (A) points at plasma membrane (PM) localization of Atg24 (see text). ll – long linker. sl – short linker. PM^Gpa2^ – the PM-targeting acylation signal of Gpa2. TMD^Sec71^ – the ER cytosol-facing targeting transmembrane domain of Sec71. Scale bar 1μm. (C) Loss of Atg24, or spatial constriction by decoration of the nanoparticle-forming Lumazine synthase of *Aquifex aeolicus* (AaLS), correlates with loss of rim opening upon partial depletion of Atg2-AID*. Atg2-AID* cells expressing mNG-Atg8 and indicated Atg24 allele were grown to logarithmic phase and treated with rapamycin, PMSF, and 10μM IAA for 3-3.5h, before imaging by widefield microscopy. Scale bar 1μm. (D) The rim-opening activity of Atg24-sfCh3V is augmented compared with Atg24-mScarletI even in unperturbed *ATG2* wildtype conditions – depending on rim recruitment by Atg20/Snx41. Cells of indicated genotype expressing mNG-Atg8 and indicated allele of Atg24 were grown to logarithmic phase, treated for 2-2.5h with rapamycin and PMSF, and imaged by widefield microscopy. Scale bar 1μm. (E) Schematic representation – the spatial extension of local Atg24C assembly is required and sufficient to open the IM rim.

To challenge this idea, we asked whether spatial constriction of Atg24C would bear a detrimental effect on rim aperture. The plasma membrane (PM) targeting acylation signal of G-protein subunit alpha 2 (PM^Gpa2^) was previously employed for PM tethering and autophagic partial loss of function of Atg17 (Suzuki *et al*., 2007). Intriguingly, PM tethering of Atg24-sfCh3V constricted not only its association with the rim but the rim itself (Figure 5A), yet this variant nevertheless mislocalized upon combined loss of Atg20 and Snx41 (Figure S5A) – suggesting active constriction of the rim by a spatially-confined Atg24C. Considering the ER association of the IM rim, which may be dysregulated upon PM tethering of the rim-associated Atg24C, we further looked for an organelle-unbiased control for the effect of lost spatial extension of Atg24C on rim aperture. Lumazine synthase of *Aquifex aeolicus* (AaLS) is a genetically encoded multimeric (GEM) nanoparticle ∼40nm-wide (Zhang et al., 2001). Strikingly, spatial confinement of AaLS-fused Atg24-mScarletI correlated with rim constriction, and a similar loss of rim visibility was observed upon complete loss of Atg24 under widefield fluorescence microscopy (Figure 5C) and CLEM (Figure 4D). The spatial extension of the autophagic Atg24C assembly is therefore tightly coupled to rim aperture upon limiting Atg2 activity.

Notably, the rim-augmenting effect of sfCh3V *versus* mScarletI fusion to Atg24 (Figure S5B) and Atg20 (Figure S5C) was observed even in mock-treated Atg2-AID* cells, suggesting augmented spatial extension of Atg24-Atg20 complex at the rim may drive opening of the rim in physiologically-relevant, Atg2-complete conditions. Accordingly, in wildtype ATG2 conditions augmented rim opening and spatially-extended assembly were observed for Atg24 (in an Atg20/Snx41-dependent manner) (Figure 5D) and of Atg20 (Figure S5D) upon fusion with sfCh3V. These observations are Vps38-independent, Atg2-dependent (Figure S5E), and independent of the w303 background, as observed in Atg11-deficient cells of the BY4741 (Figure S5F) or SEY6210 (Figure S5G) background. Spatial extension of the Atg24 complex assembly is thus required and sufficient to open the autophagic membrane rim (Figure 5E), thus opposing the constriction promoted by the Atg2-Atg18 complex (Figure 2).

### Excessive activity of the Atg24-Atg20 complex impairs IM maturation

Following the visible widening effect of the Atg24 complex on the rim, we asked whether excessive Atg24 activity might impair vesicle maturation. First, we noted that mild depletion of Atg2 led to an impaired flux of GFP-Atg8 (Figure S2A) but not due to lost membrane formation and initial expansion (Figure 2A), suggesting that loss of flux occurred at a later step. On the other hand, as less autophagic bodies accumulate over time upon titration of Atg2-AID* by IAA, it follows that autophagy was impaired prior to vacuolar fusion (Figure S6A), at either the rate of expansion, the rate of maturation – or both. The ascribed role of Atg2 in IM expansion (Noda, 2021) would suggest a reduced rate of expansion upon partial loss of Atg2. Nevertheless, we asked whether subsequent sealing or clearance of associated factors are also compromised. Notably, partial depletion of Atg2 largely maintains the size of rapamycin-induced IMs (Figure S6B) and overnight nitrogen starvation of partially-depleted Atg18-AID* cells showed impaired vacuolar delivery of Atg8, accompanied by retention of an abnormally-large IM (Figure S6C). These observations suggest that partial loss of Atg2-Atg18 activity may delay sealing even following sufficient membrane expansion.

The stable association of Atg24 with the excessively-wide rim (Figure S3F) hints at a possible role for Atg24 complex in impairment. Notably, spontaneous dislodging of Atg24 from the Atg2-depleted rim was accompanied by rim constriction and loss of Atg8 fluorescence (Figure 6A), suggesting that alleviation of an inhibitory role of Atg24C promotes maturation. Moreover, sfCh3V-fused Atg24, which was shown to confer excessive opening of the rim (Figure 5), also impaired accumulation of vacuolar autophagic bodies (ABs) under conditions of partial Atg2 depletion (Figure S6D), in line with an inhibitory role of Atg24C in maturation. This is also indicated by western blot analysis observation that the flux of GFP-Atg8 was more severely impaired by partial depletion of Atg2-AID* upon fusion of Atg24 to sfCh3V compared with mScarletI (Figure S6E). Accordingly, under Atg2-limiting conditions, Atg24-sfCh3V stably associates with the rim (Figure S6F), but dislodges from the rim upon closure (Figure 6B). We then asked whether artificially-excessive Atg24-Atg20 activity may compromise sealing in wildtype Atg2 conditions. Strikingly, in Vps38-deficient cells the flux of GFP-Atg8 compared is severely compromised by fusion of Atg24 to sfCh3V compared with mScarletI in presence of comparable Atg2 levels (Figure 6C). Moreover, Atg20-sfCh3V dislodges from the excessively-open rim rims upon closure in wildtype cells (Figure 6D). Taken together, these data indicate that Atg24C has an intrinsic capacity to impair IM maturation and flux upon excessive widening of the rim.

**Figure 6:**
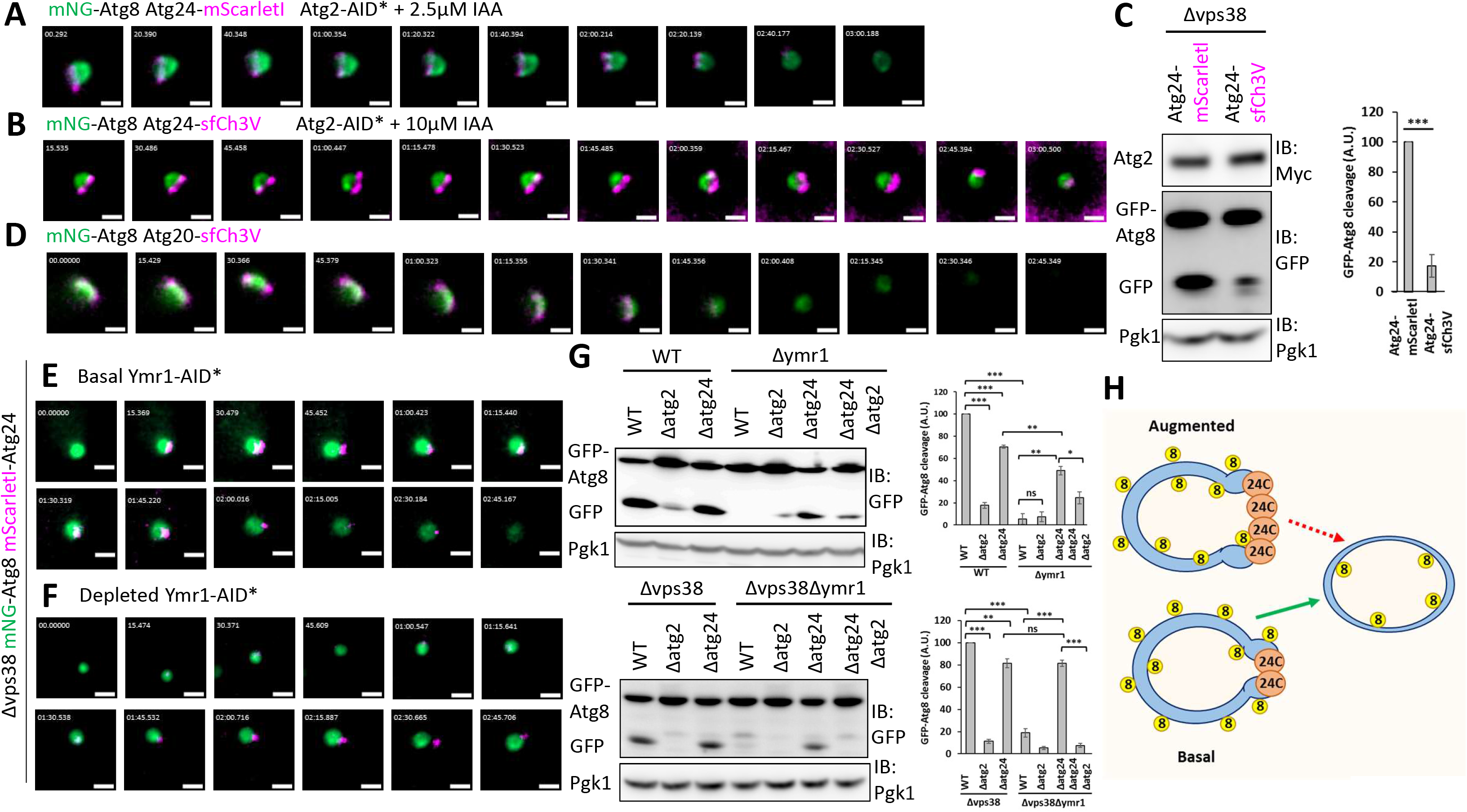
Excessive activity of the Atg24 complex impairs IM maturation. (A, B, D) Atg24 and Atg20 dislodge from the rim concomitantly with IM maturation. Atg2-AID* (A, B) or wildtype (D) cells expressing mNG-Atg8 and Atg24-mScarletI (A), Atg24-sfCh3V (B), or Atg20-sfCh3V (D) were grown to logarithmic phase, treated for 1.5-3.5h with rapamycin, as well as 2.5μM IAA (A) or 10μM IAA (B) and imaged by widefield microscopy at indicated time-points. Scale bar 1μm. (C) The rim-augmenting activity of Atg24-sfCh3V impairs the autophagic flux of GFP-Atg8 – in comparison with Atg24-mScarletI – in absence of Vps38. Δvps38 cells expressing Atg2-8xMyc, GFP-Atg8, and indicated Atg24 allele were grown to logarithmic phase, treated for 3h with rapamycin and extracted proteins were immunoblotted as indicated. Error bars – S.E.M from 3 experiments, paired t-test: *** p<0.0001. (E, F) Depletion of Ymr1-AID* leads to retention of Atg24 at the IM rim – in absence of Vps38. Δvps38 Ymr1-AID* cells expressing mNG-Atg8 and mScarletI-Atg24 were grown to logarithmic phase, treated for 1-2h with rapamycin and mock (0μM) or saturating (250μM) IAA for indicated basal or depleted Ymr1 levels, respectively, and imaged by widefield microscopy at indicated time-points. Scale bar 1μm. (G) Loss of Atg2-dependent flux of GFP-Atg8 in absence of Ymr1 is partially suppressed by further loss of Atg24 – in a Vps38-independent manner. Cells of indicated genotype expressing GFP-Atg8 were grown to logarithmic phase, treated for 4h with rapamycin and extracted proteins were immunoblotted as indicated. Error bars – S.E.M from 3 experiments, paired t-test: * p<0.05, ** p<0.001, *** p<0.0001. (H) Schematic representation – an excessive opening of the IM rim by augmented Atg24C activity compromises maturation.

Finally, we asked whether the endogenous unperturbed Atg24 protein may also play an inhibitory role in maturation. The PI(3)P phosphatase Ymr1 has been implicated in maturation by turnover of PI(3)P and consequent clearance of associated factors (Cebollero *et al*., 2012). As Atg24 requires PI(3)P for association with membranes, we wondered whether unchecked retention of Atg24 at the rim of Ymr1-deficient IMs may underlie their maturation defect. Upon mock-depleted, basal levels of Ymr1-AID* in Vps38-deficient cells, Atg24 dynamically associates with the rim but leaves at maturation (Figure 6E). Transient depletion of Ymr1-AID*, however, leads to retention of Atg24 at the rim of the expanded, maturation-defective expanded isolation membranes (Figure 6F), suggesting an inhibitory role for Atg24C in maturation. Accordingly, western blot analysis indicates that Atg2-dependent autophagic flux is lost upon Ymr1 knock-out, but rescued by further loss of Atg24 – in a Vps38-independent manner (Figure 6G). Taken together, our observations link excessive, dysregulated activity of the Atg24 complex at the rim to compromised maturation and consequently impaired autophagic flux (Figure 6H).

### Atg24 couples IM maturation with sequestration of large cargo

The Atg24-Atg20/Snx41 complex has been previously implicated in starvation-induced autophagy of fatty acid synthase (FAS) (Shpilka et al., 2015), mRNA (Makino et al., 2021) and the 26S proteasome, as well as the small (40S) and large (60S) subunits of the 80S eukaryotic ribosome (Nemec et al., 2017) – which is ∼30nm-wide (Ben-Shem et al., 2011). Still, the underlying mechanism remained unclear thus far, considering that a wide-open rim is expected to accommodate different cargos regardless of their size. In light of the narrow rim of the wildtype isolation membrane compared with the large vesicle body in our hands (Figure 1), and considering the *in situ* autophagy-specific role of the Atg24 complex in its opening (Figure 5), we checked whether Atg24 may play a physiological role in sequestration of large cargo. As negative controls, the small cytosolic enzymes Fba1 and Pgk1 appeared in ABs regardless of Atg24 depletion (Figure 7A). Strikingly, the Rps1b and Rpl9b subunits of the small (40S) and large (60S) ribosomal particles, respectively, were observed in autophagic bodies in an Atg24-dependent manner (Figure 7A), in line with the CLEM observation of ribosome exclusion from the interior of Atg24-deficient IMs (Figure 4D).

**Figure 7:**
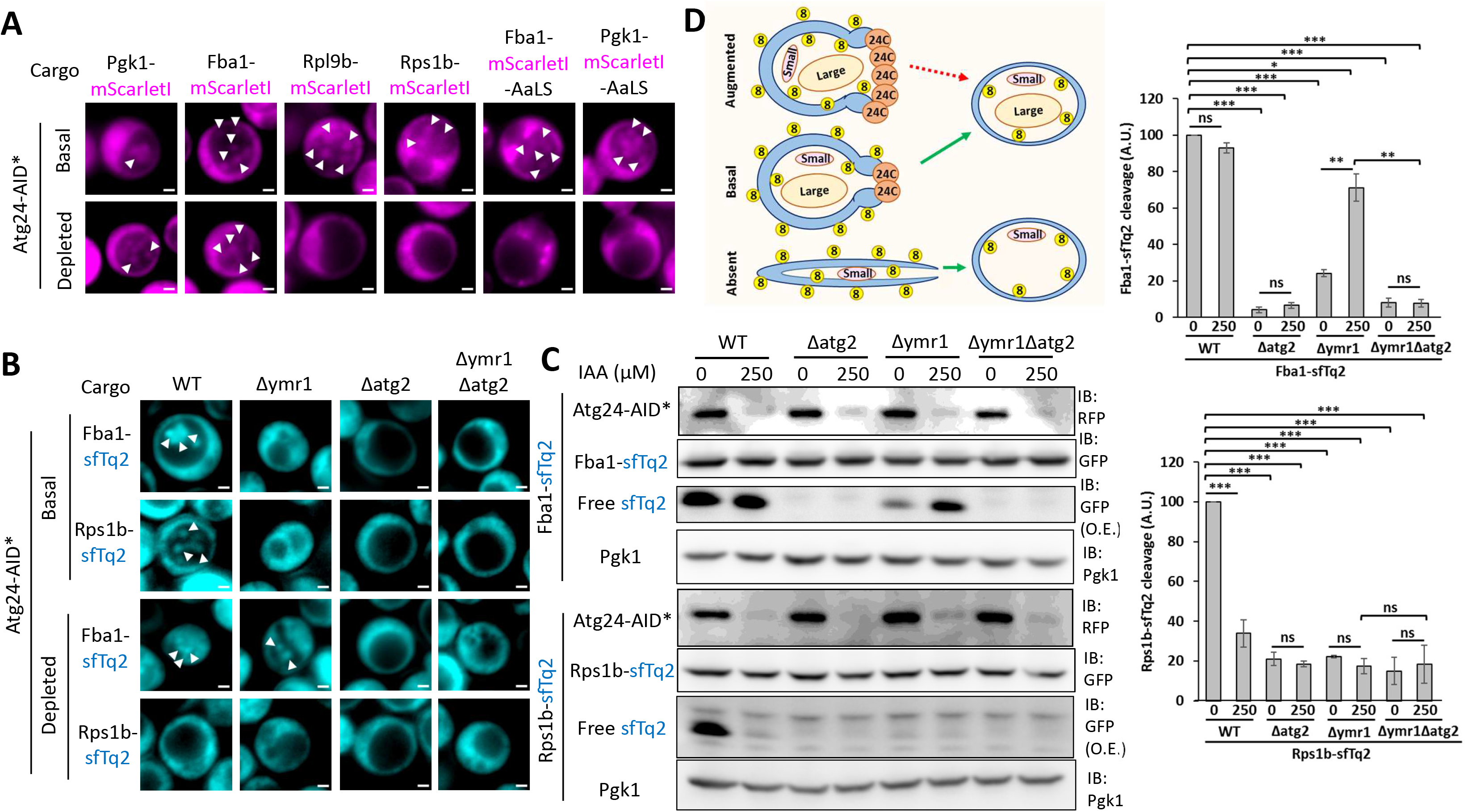
Atg24 couples IM maturation with sequestration of large cargo. (A) Atg24 is dispensable for formation of autophagic bodies including small cargo proteins Pgk1 and Fba1 – but essential for autophagy of large cargo proteins 40S ribosomal subunit Rps1b, 60S ribosomal subunit Rpl9b and the Pgk1/Fba1-decorated the *Aquifex aeolicus* lumazine synthase (AaLS) nanoparticle. Atg24-AID* cells expressing indicated mScarletI-tagged cargo protein were grown to logarithmic phase, treated for 5h with rapamycin, PMSF, and mock (0μM) or saturating (250μM) IAA for indicated basal or depleted Atg24 levels, respectively, and imaged by widefield microscopy. Arrowheads point at autophagic bodies. Scale bar 1μm. (B, C) The loss of Atg2-dependent vacuolar delivery of cargo in absence of Ymr1 can be partially suppressed by further loss of Atg24 – for small cargo Fba1 but not large cargo Rps1b. Atg24-AID*-sfCh3V cells with indicated genotype expressing mNG-Atg8 and sfTq2-fused indicated cargo protein were grown to logarithmic phase, and treated for 4-5h with rapamycin, mock (0μM) or saturating (250μM) IAA for indicated basal or depleted Atg24 levels, respectively, and PMSF (B) before imaging by widefield microscopy (B) immunoblotting or extracted proteins as indicated (C). Arrowheads point at autophagic bodies. Scale bar 1μm. Error bars – S.E.M from 3 experiments, paired t-test: * p<0.05, ** p<0.001, *** p<0.0001. (D) Schematic representation – fine-tuning of rim-localized Atg24 complex activity may balance autophagic sequestration of large cargo at the course of expansion on one hand with eventual maturation of the autophagic vesicle on the other hand.

To exclude putative cargo-specific effects of Atg24 (Henkel et al., 2020; Waite et al., 2022), we engineered artificially-large cargos from small endogenous proteins by decorating the ∼40nm-wide AaLS nanoparticle, previously employed in yeast for its size-induced slower diffusion (Delarue et al., 2018), with Fba1 or Pgk1. Like the 40S and 60S ribosomal proteins examined above, Fba1 or Pgk1-decordated AaLS appeared in ABs in basal but not depleted Atg24 conditions (Figure 7A). Importantly, these observations were Atg2-depndent but Vps38-independent (Figure S7A) – indicating an autophagy-specific, endosomal-independent role of the Atg24 in cargo sequestration. Moreover, western blot analysis of autophagic flux, indicated by of vacuolar cleavage of sfTq2-fused cargo, showed loss of Atg2-dependent flux for Rps1b but not Fba1 upon transient depletion of Atg24-AID* (Figure 7C). Importantly, vacuolar delivery of large cargo (40S subunit Rps1b or 60S subunit Rpl8a) required Atg24 even upon forced interaction of cargo with Atg8, tagged by GFP-derived sfTq2, by fusion of cargo to a GFP-binding protein (GBP). As controls, GBP-fused small cargo Fba1 – as well as sfTq2-Atg8 itself – appeared in ABs regardless of Atg24 (Figure S7B). Taken together, these observations indicate that Atg24 is essential for entry of large – cargo – but not of small cargo – into the strictly-nonselective isolation membrane. Out data is in line with *passive* sequestration of large cargo that is likely compromised *physically* due to loss of rim opening activity of Atg24 (Figure 5C), rather than *active* recruitment of specific cargo by an Atg24-mediated pathway (Nemec *et al*., 2017) that is *kinetically* impaired in its absence.

Finally, as loss of Atg24 not only compromises sequestration of large cargo but also suppress the maturation defect of Ymr1-deficiency (Figure 6G), we wondered whether these physiological and mechanistic effects, respectively, are coupled. For this, we analyzed by widefield microscopy entry of cargo into vacuolar ABs (Figure 7B), coupled with western blot analysis of vacuolar cleavage of sfTq2-tagged cargo (Figure 7C). Both assays demonstrate that the impairment in Atg2-dependent autophagic flux due to loss of Ymr1 may be suppressed for the by further depletion of Atg24 for the small cargo Fba1 – but not for large cargo subunit Rps1b. Microscopy of Atg8 and Atg24 under these conditions (Figure S7C) confirms the established Atg2-dependent rim association of Atg24 (Figure 3) and the morphological effect of its depletion (Figure 4) under these conditions. Overall, our observations indicate that loss of Atg24 may rescue vesicle maturation and autophagic flux for small cargo in absence of Ymr1, in the cost of compromised autophagic inclusion of large cargo in presence of Ymr1. We therefore surmise that fine spatiotemporal tuning of rim stabilizing and opening activity of the PI(3)P effector complex Atg24-Atg20/Snx41, e.g. by the PI(3)P phosphatase Ymr1, *versus* the rim-constricting activity of the antagonistic PI(3)P effector complex Atg2-Atg18, allows autophagic sequestration of large cargo in the course of IM expansion on one hand, while securing eventual maturation of the autophagic vesicle on the other hand (Figure 7D).

## Discussion

In selective autophagy cup-shaped phagophores expand in spatiotemporal coordination with tight engulfment of cargo (Schutter *et al*., 2020; Suzuki *et al*., 2013), and mature upon completion of engulfment. In nonselective autophagy, however, coordination of these events in context of the prevailing cup-shaped paradigm of IM expansion has been elusive. Here we show that *bona fide* nonselective IMs expand in a spherical shape while maintaining a narrow rim (Figure S7D, “Basal”), reminiscent of amphoras – ancient storage vessels of a large body with a narrow rim curved outwards. We therefore term these strictly non-selective IMs “amphores”, distinguished from strictly-selective “phagophores”. This distinction fits recent detection of mammalian amphora-shaped IMs containing strictly cytosol, compared with apparently wider rims of IMs that engulf fragments of organelles (Takahashi et al., 2022). The amphore model is substantiated by our molecular characterization of opposing forces that govern rim aperture *in situ* – constriction by the Atg2-Atg18 complex and widening by the autophagy-specific activity of Atg24 that assembles at the rim with Atg20 or Snx41 upon autophagy-specific PI(3)P formation. The topology of an amphore with a narrow rim rather than a wide-open phagophore explains why rim widening by Atg24 is dispensable for vesicle expansion and sequestration of small cargo, yet essential for sequestration of large cargo (Figure S7D, “Absent”).

The morphogenic effect of Atg24 on IM shape (Figure S7D, “Absent”) suggests that sequestration of large cargo scaffolds the expanding vesicle. Actin has been implicated in scaffolding the mammalian autophagosome (Zientara-Rytter and Subramani, 2016) but is dispensable for nonselective autophagy in yeast (Reggiori et al., 2005). Alternatively, the Atg12∼Atg5-Atg16 complex has been implicated in scaffolding the yeast phagophore (Kaufmann et al., 2014), in line with the association of Atg5 and Atg16 with the selective cargo surface (Suzuki *et al*., 2013). However, in nonselective autophagy Atg5 visibly localizes only to the vacuolar membrane (VM)-associated base of the IM (Graef *et al*., 2013) (Figure S1A). Here we speculate that bulk mechanical pressure on the growing membrane by collision with large cytosolic particles like ribosomes is countered by outward-facing pressure of large sequestered cargo in presence of Atg24 – but flattens the membrane under Atg24-deficient conditions in its absence. The amphore topology also accounts for impairment in maturation upon excess widening of the rim by the Atg24 complex – either spatially due to augmented assembly (Figure S7D, “Augmented”), or temporally due to loss of PI(3)P turnover by Ymr1. As the loss of Ymr1-dependent autophagic flux is completely recovered by further elimination of the autophagic activity of Atg24 (Figure 6G, compare Δatg24 with Δymr1Δatg24 under Δvps38), we surmise that PI(3)P turnover facilitates maturation by primarily by of Atg24 assembly from the rim to facilitate maturation. Taken as a whole, our experimental analysis of the amphore model suggests that fine-tuned widening of the nonselective IM rim balances size-dependent cargo sequestration at the course of vesicle expansion with productive maturation for size-independent cargo consumption. The amphore model thus ties together the topology, molecular mechanism, and physiology of nonselective autophagy.

The low local abundance of IM-associated autophagic proteins (Geng *et al*., 2008) suggests that a narrow IM rim is more favorable than a wider rim for quantitative regulation by Atg2. Previous studies of Atg2 indicated a structural role in bridging the phagophore rim with an endoplasmic reticulum (ER) domain (Gomez-Sanchez *et al*., 2018; Kotani *et al*., 2018) and an *in vitro* lipid transport activity (Osawa *et al*., 2019) – both depending on its N-terminal Chorein domain. Intriguingly, an experimentally-validated Atg1 phosphorylation site immediately follows this domain (Hu et al., 2019), whose loss may be partially compensated by its homolog in the multiple contact site lipid transporter Vps13 (Osawa *et al*., 2019). Maintenance of membrane growth upon partial loss of Atg2 activity in our hands is in line with the recently-reported compensation for down-regulation of Atg2 by the local activity of Vps13 at the rim (Dabrowski et al., 2022). The maintenance and augmentation of Atg24 rim activity upon ER tethering in our hands (Figure 5A) calls for characterization of the ER domain where Atg2 and Atg24 activities converge. This yeast-specific domain may be the functional analog to the mammalian omegasome ER domain marked by DFCP1 (Axe et al., 2008), a FYVE (Fab1, YGL023, Vps27, and EEA1) domain protein which binds PI(3)P (Derubeis et al., 2000) – like Atg24.

The requirement for a binding partner of Atg24 in rim targeting (Figure 3B) is compatible with reported motifs in the BAR (Bin/Amphiphysin/Rvs) domain of Atg20 that govern its interaction with the BAR domain of Atg24 (Popelka et al., 2017), recently shown as essential for membrane binding and remodeling *in vitro* (Reinhart *et al*., 2022). Future mutational studies are required to correlate augmented *in vitro* membrane remodeling activity with the competence to stabilize and promote rim opening. The rim of autophagic membranes appears to be deformed locally (Figure 2B) (Bieber *et al*., 2022), which could explain the contribution of the membrane curvature-binding BAR domain (Peter *et al*., 2004) of Atg24 to rim targeting (Figure 3A). Excessive association of Atg24 and Atg20 with the wide-open rim upon partial loss of Atg2 (Figure 3D, 3E) could thus be attributed to a marked increase in surface area of the destabilized curved rim membrane. Moreover, as compromised expression of Atg2 in *Drosophila melanogaster* leads to aberrant accumulation of phosphatidylserine (PS) in autophagic membranes (Laczko-Dobos et al., 2021), favored recruitment of the Atg20-Atg24 complex by elevation in PS (Ma et al., 2018) may underlie this response of the Atg24 complex to loss of Atg2. Strikingly, we recently correlated a sharp increase in PS content with abnormal retention of unsealed, wide-open autophagic membranes upon loss of the yeast phosphatidylcholine synthesis enzyme Opi3 (Polyansky et al., 2022) – suggesting a shared mechanism for these phenotypes.

In the context of nonselective, Atg11-independent autophagy we observe that the Atg24 complex stabilizes a wide rim in general and Atg20 is capable of promoting excessive widening of the rim in Atg2-complete conditions in particular (Figure S5D). These observations raise the possibility that the Atg24-Atg20 complex may directly promote substantial widening of the IM rim during Atg11-mediated selective engulfment of large cargo. Atg24 and Atg20 were originally localized to the selective, Ape1-associated IM (Nice *et al*., 2002), and both are genetically required for pexophagy (Nice *et al*., 2002) and mitophagy (Okamoto et al., 2009), and their fission yeast homologs promote organelle autophagy (Zhao et al., 2016). However, the contribution of the Atg24-Atg20/Snx41 complex to selective autophagy was later attributed to trafficking of Atg9, in starvation-induced redundancy with other endosomal retrograde pathways (Ohashi and Munro, 2010). While Snx41 is implicated in retrograde trafficking of the Atg9-vesicle biogenesis factor Atg27 (Ma *et al*., 2017), Atg20 directly interacts with Atg11 for a yet-unknown function (Popelka *et al*., 2017). It would be therefore be interesting to test whether the Atg24-Atg20 complex localizes to the leading edge of selective IMs to facilitate cargo engulfment. Based on our observations from nonselective autophagy, it may be predicted that artificial augmentation of their rim expanding capacity would contribute to the rate of the process or to its efficacy for oversized cargos that otherwise fail to be engulfed.

Intriguingly, Atg24 is essential for vacuolar fusion in a genetic setting of endosomal – rather than mitochondrial – PS synthesis (Ma *et al*., 2018). Moreover, in the aging model organism *Podospora anserina*, Atg24 is only partially required for selective autophagy but is necessary for efficient nonselective autophagy (Henkel *et al*., 2020) – as is the case for the white head blight pathogen *Fusarium graminearum* Atg20 (Lv et al., 2020). These studies indicate that under certain physiological conditions or genetic settings the Atg24 complex may support sufficient opening of the rim even for sequestration of small cargo on one hand while extending to rim-independent, endosome-associated roles on the other hand. The reported interactions of the nonselective autophagy scaffold Atg17 with Atg24 (Nice *et al*., 2002) and with Atg11 (Yorimitsu and Klionsky, 2005) asks for elucidation of the manner by which Atg11 and Atg17 coordinate assembly of the Atg24 complex on selective, nonselective and putatively mixed selectivity IMs. Furthermore, given the diverging roles of Atg24-Atg20 and Atg24-Snx41 complexes in endosomal trafficking (Ma *et al*., 2017), partitioning of the Atg24 complex between the endosomal and autophagic pathways, by differential regulation of their respective PI(3)P signaling activities, should be quantitatively linked to the balance between sequestration of large cargo and maturation of the autophagic vesicle.

## Acknowledgments

Z.E. is the incumbent of the Harold Korda Chair of Biology. We are grateful for funding from the Israel Science Foundation (Grant #215/19), Joint NRF - ISF Research Fund (Grant #3221/19), Joint NSFC-ISF Research Fund (Grant # 3345/20), and the Yeda-Sela Center for Basic Research. We thank Olena Trofimyuk and other members of the Elazar group for their experimental support. Leica HyVolution confocal and STED images were acquired at the Advanced Optical Imaging Unit, de Picciotto-Lesser Cell Observatory unit at the Moross Integrated Cancer Center Life Science Core Facilities, The Weizmann Institute of Science.

## Author contributions

O.S. – Conceptualization, Methodology, Investigation, Data Curation, Writing – Original Draft, Writing – Review & Editing, Visualization

M.F. – Investigation, Data Curation, Formal Analysis, Writing – Review & Editing

A.P. – Investigation, Data Curation

E.S. – Investigation, Visualization

T.D. – Investigation, Visualization

S.W. – Investigation, Visualization, Data Curation

Z.E. – Conceptualization, Supervision, Writing – Original Draft, Writing – Review & Editing, Project Administration, Funding Acquisition

## Methods

### Yeast strains

Standard yeast maintenance, growth and transformation methods were used. Strains used in this study and details pertaining to their generation and experimental application are listed Table 1, and were generated using oligonucleotides (Sigma) listed in Table 2 and plasmids listed in Table 3. All plasmids were taken from our lab stock and carry a yeast selection marker and an AmpR *E. coli* resistance cassette on a pFA6 backbone (Wach et al., 1994), with the following exceptions. pOS7 (kind gift from Bernd Bukau) was published as pNHK53 (Nishimura *et al*., 2009) with undocumented backbone. pOS202 (kind gift from Maya Schuldiner) is derived from pTIR2 (Nishimura *et al*., 2009) and was integrated seamlessly without a selection marker (see ORS475 below). pFA6-kanMX4 (Wach *et al*., 1994) and pFA6-natMX4, pFA6-hphMX4 (Goldstein and McCusker, 1999) were published. pOS642 (kind gift from Michael Thumm) was published as vac-BFP (Graef *et al*., 2013) and derives from pRS305. The TAP (tandem affinity purification) tag in our plasmids contains 6xHis-3xFLAG-3xFLAG-TwinStrepTag.

All strains were derived by a series of stable chromosomal integrations onto a w303 wildtype strain (kind gift from Jeffrey Gerst), except where SEY6210 (kind gift from Jeffrey Gerst) or BY4741 (from our lab stock) backgrounds are indicated. All strains employed in experiments express exogenous Atg8, tagged at its N-terminus with mNG, sfTq2 or yeast-enhanced GFP (Meiresonne et al., 2019; Shaner et al., 2013), under its native promoter and terminator, integrated in place of the ATG11 open reading frame (ORF) using ORF flanking homologous primers that anneal at the ATG8 promoter and downstream of the hphNT1 cassette. Endogenous Sec61 was visualized by C-terminal tagging with HaloTag7 (Ohana et al., 2009). Pho8 was visualized by exogenous expression of an additional copy of Pho8 tagged at its N-terminal with TagBFP (Subach et al., 2008) under its native terminator and the PGK1 promoter. PI(3)P was visualized by exogenous expression under the pCUP1 promoter of the PX domain of Vam7 (PX^Vam7^), tagged with mScarletI (Bindels et al., 2017). The *delitto perfetto* two-step scarless procedure (Storici and Resnick, 2006) was employed for generation of ORS475 from ORS345 *via* ORS474. Linearization of auxotrophic marker integration plasmids by treatment of plasmids pOS7 and pOS642 with the restriction enzymes StuI and BstII (NEB) were transformed for generation of ORS1150 and ORS1787, respectively.

Otherwise, all strains were generated by the S1/2/3/4 homologous recombination PCR integration method (Janke et al., 2004) using a Q5 PCR Kit (NEB). S1-S2 primers was used to knock out endogenous proteins, with optional knock-in of the exogenous mScarletI-PX^Vam7^ expression cassette, S3-S2 primers were used for C-terminal tagging of endogenous proteins, expressed under their native promoter, and S1-S4 primers were used for N-terminal tagging of endogenous proteins expressed under the pCUP1 promoter or for down-regulation by promoter replacement with the pCUP1(dim) promoter – as indicated. After transformation and isolation of a single colony grown on selective plates according to indicated yeast marker, integration of the insert into the expected genomic locus was verified by colony PCR using HyTAQ ReadyMix (HyLabs), reverse primer EW_NATR (except where noted) annealing inside the insert, and forward primer annealing upstream of the integrated locus – as indicated in Table 1.

### Experimental procedures

All experiments were performed at 30C (in 180-220rpm agitation, except for live microscopy). Yeast were inoculated from a patch on a YPAD plate into SD medium: 0.19g/l LoFlo YNB without nitrogen (CYN6201, Formedium), 5g/l ammonium sulfate (Merck), 2% (w/v) glucose (Sigma) and amino acid mix (100mg/L glutamic acid, 35mg/L histidine, 110mg/L leucine, 120mg/L lysine, 40mg/L methionine, 50mg/L phenylalanine, 375mg/L serine, 200mg/L threonine, 50mg/L tryptophan, 40mg/L uracil, 50mg/L adenine). After overnight growth as above to saturation, yeast were diluted and grown to logarithmic phase. Expression of mScarletI-PX^Vam7^ from the pCUP1 promoter or of down-regulated proteins from the weak pCUP1(dim) promoter was induced by supplementation of 10μM (unless noted otherwise) copper sulfate (Merck) to the growth medium, while other proteins tagged at their N-terminus and driven by pCUP1 promoter were expressed at basal promoter activity by the copper composition of the growth medium. HaloTag7-fused proteins were visualized by centrifuging 1ml culture (1000g, 2min, RT) as above, resuspending in 20-100μl same culture containing 100nM Halo-JF635 (Grimm et al., 2017) from 2-10μM stock in DMSO and incubated at least 30 minutes prior to dilution (at least 1:10) into stain-free medium. Proteins were depleted where indicated by chromosomally tagging the C-terminus of the coding sequence to the minimal *Arabidopsis thaliana* IAA17 auxin-inducible degron (AID*) domain (aa 71-114) and addition of 0μM (mock depletion), 1-10μM (partial depletion) or 250μM (saturated depletion) indole-3-acetic acid (IAA) from 200x stock in a 50:50% (v/v) DMSO:EtOH solvent to the culture 0-30mins before induction of autophagy.

Autophagy was induced by treatment with 400ng/ml rapamycin (GA7417, Glentham) from 400ug/ml in DMSO, accompanied (except where indicated) with inhibition of vacuolar processing of autophagic bodies by 1mM PMSF (P7626, Sigma) from 200mM stock in 100% (v/v) EtOH. For western blot analysis or correlative light electron microscopy (CLEM) cells were treated directly in a 1ml microtube, while for widefield fluorescence or Leica HyVolution microscopy 5-10μl culture was diluted into 90-95μl induction medium within 384-well plates (P384-1.5H-N, CellVis), optionally pre-coated at RT for 20min with 20μl of 0.2mg/ml in DDW concanavalin A (L7647, Sigma) and air-dried.

### Western blot analysis

1ml treated cell culture (0.7-1 OD 600) was optionally treated by 10% (v:v) TCA and further neutralized by 10% unbuffered Tris, and pellets were frozen in liquid nitrogen and stored at -80C. After thawing on ice, pellet was treated with 0.1M NaOH (10min, ice), centrifuged (1000g, 2min), resuspended in Laemmli sample buffer (100 µl 1x), and heated (95°C, 5 min). Proteins were analyzed on 4-12% or 8-16% SDS-PAGE gradient gels (Invitrogen) and transferred to nitrocellulose membrane (BioRad). Membranes were blocked (5% w:v non-fat milk powder in PBS, 1h), incubated with indicated primary antibody (2h RT, or overnight at 4C), washed 3×5-10min PBST (PBS + 0.1% (v:v) Tween-20), incubated with HRP-conjugated secondary antibody (in 2.5% (w:v) non-fat milk powder in PBST, 1h), washed 3xPBST and visualized using the EZ-ECL western blotting detection reagent (Biological Industries). Protein band intensity was quantified in ImageJ (NIH). Vacuolar processing of GFP-Atg8 into free GFP (Meiling-Wesse et al., 2002) or of cargo fused to the GFP derivative sfTq2 (Welter et al., 2010), was calculate in Microsoft Excel by dividing the amount of the free fluorescent protein (FP) from the sum of free FP and cargo-fused FP.

### Widefield fluorescence microscopy

Plate was mounted with immersion oil (Immersol 518F 30C, Zeiss) in widefield microscopy (AxioObserver7, Zeiss) on objective (Plan Apochromat 100x/1.46 NA, Zeiss) in cage pre-heated to 30C. Microscope was operated by computer software (ZEN Blue 3.2, Zeiss) to capture image z-stacks (4-5.76μm deep, 0.24-0.40μm interval) by an ORCA Flash 4.0 v3 camera (Hamamatsu) through a 1.6x tube lens, illuminated by bright field illumination with DIC optical path (NA 0.55) or by LED illumination (Colibri 7, Zeiss). For mNG, mScarletI, (TagBFP, Halo-JF645) imaging, excitation LEDs were 475nm, 555nm (, 385nm, 630nm), respectively, and the 90 HE filter set (Zeiss) was used. For mNG/GFP, mScarletI/sfCh3V imaging, excitation LEDs were 475nm, 590nm, respectively, and the 92 HE filter set (Zeiss) was used. For sfTq2, mNG (, sfCh3V) imaging, excitation LEDs were 430nm, 511nm (, 590nm), respectively, and the 91 HE filter set (Zeiss) was used. Images were smoothed by a Gaussian filter (radius 1.30), optionally deconvolved by a constrained-iterative algorithm and adjusted for display manually, for elimination of uniformly-diffuse background signal at the low threshold and allow 0.2% saturation at the high threshold.

### Leica HyVolution confocal microscopy

Confocal imaging was performed using an inverted Leica SP8 STED3X microscope, equipped with Acusto Optical Tunable Filter (Leica microsystems CMS GmbH, Germany), HCX PL APO 93x/1.30 GLYC STED White motCORR objective, and a white light laser (WLL). mNG-Atg8 was excited at 501nm, z-stack were acquired using a galvo stage with 0.05µm intervals, and emission was collected using a galvometric scanner operating at 400Hz and an internal HyD detector at the range of 511nm-537nm with line averaging 4. Pixel size and zoom were 76nm,17.3 for Δvps38Δymr1 imaged under standard confocal mode, and 43nm,15.78 for Δvps38Δymr1Δatg24 imaged under STED mode with a depletion line at 592nm. The acquired images were visualized using LASX software (Leica Application Suite XLeica microsystems CMS GmbH) and processed with the Lightning module in adaptive mode with refractive index 1.4 and default parameters.

### Correlative light electron microscopy

1ml of yeast culture was pelleted (500g, 1min), the pellet was placed in an aluminum disc with depression of 100 µm and outer diameter of 3 mm (Engineering Office M. Wohlwend GmbH, Sennwald, Switzerland) and was covered with a matching flat disc. The sandwiched sample was high-pressure frozen using an EM ICE high pressure freezing device (Leica Microsystems, Vienna Austria). The frozen samples were dehydrated by freeze substitution in an AFS2 freeze substitution device (Leica Microsystems, Vienna Austria) in anhydrous acetone containing 0.1% uranyl acetate, embedded in Lowicryl HM20 acrylic resin (Electron Microscopy Sciences, USA) and polymerized in UV according to (Kukulski et al., 2012). Sections with thickness of 500 nm were cut with a diamond knife (Diatome, Biel, Switzerland) using a UC 7 ultramicrotome (Leica Microsystems, Vienna, Austria). Sections were mounted on 200 mesh Formvar coated nickel grids. Widefield fluorescence images were taken in order to identify yeast cells expressing mNG-Atg8 and Atg24-mScarletI using VUTARA SR352 system (Bruker) with 1.3 NA 60x silicon oil immersion objective (Olympus). Z slices of 250 nm were collected using 488 nm and 561 nm excitation lasers in the presence of a buffer containing 7 μM glucose oxidase (Sigma), 56 nM catalase (Sigma), 2 mM cysteamine (Sigma), 50 mM Tris, 10 mM NaCl, 10% glucose, pH 8. Chromatic calibration of the 488 and 561 nm channels was performed prior to data collection using glass slide with tetraspeck beads (Invitrogen T7279). Images of both channels were aligned using Vutara software (Bruker). The correction was applied to the yeast images. The same grids were washed with bi-distilled water, incubated in droplet of 1 mg/ml polylysine solution for 1 minute, washed in 3 droplets of DDW, incubated in a droplet of colloidal gold (diameter ∼12 nm), and washed in one droplet of DDW. The grids were blotted and then double stained with 2% uranyl acetate and Reynolds lead citrate. The grids were then coated with ∼2nm carbon using a CCU-010 carbon coater (Safematic, Switzerland). Sections were viewed using a Tecnai TF20 transmission electron microscope (Thermo Fisher Scientific, Eindhoven, the Netherlands) operating at 200 kV using the STEM mode. The SerialEM program (Mastronarde, 2005) was used for acquisition of large regions for orientation and for correlating between images acquired by light microscopy and regions observed in the STEM as well as for the automated acquisition of STEM tomograms. Initial registration between LM and TEM regions was conducted using the grid’s mesh corners as registration points. mNG and mScarletI labeled regions were used to identify relevant yeast cells in the section and regions of interest for collecting STEM tomograms. Fine-tuning of the correlation was based on cell shape and relation to neighboring cells. Automated STEM tomography datasets of targeted regions were collected at the Tecnai TF20 microscope, using SerialEM software. Images were recorded every 1° between -60° to 60°, using a Fischione HAADF detector set up to perform as a Bright-Field detector, by inserting an objective aperture and positioning the beam over the HAADF detector. The 2K by 2K images were taken at 40,000 magnification, corresponding to a pixel spacing of 1.4 nm. Tomograms were reconstructed using the IMOD software suite (Kremer et al., 1996). Overlay of the chromatic corrected fluorescence images (of both 488 and 561 nm channels) and the tomography data was performed using Adobe Photoshop software. For each cell the best fluorescence z slice was selected and overlaid with the corresponded tomography virtual slice.

### Statistical analysis

Graph bars represent standard error of the mean (S.E.M) of indicated number of experiments. Statistical significance of difference between indicated averages was calculated in GraphPad by a paired two-tailed t-test: ∗, p < 0.05; ∗∗, p < 0.01; ∗∗∗, p < 0.001, ns – non-significant.

## Supplemental Information

Supplemental Table 1: Yeast strains used in this study

Supplemental Table 2: Oligonucleotides used in this study

Supplemental Table 3: Plasmids used in this study

Movie M1: Corresponding to Figure 2B

Movie M2: Corresponding to Figure 4D

