## Supplementary figures and text for "Rim aperture of autophagic membranes balances cargo inclusion with vesicle maturation"

### Supplemental Figure Legend

#### Figure S1: Nonselective isolation membranes (IMs) expand as large vesicles with a narrow rim

(A) Canonical association pattern of autophagy proteins with the nonselective isolation membrane (IM) – Atg1, Atg2, and Atg5 localize throughout the IM surface, at its base, and at its rim, respectively. Wildtype or Δvps38Δymr1 cells as indicated expressing sfTq2-Atg8 and indicated mNG-tagged autophagy factors were grown to logarithmic phase, treated for 2-3.5h with rapamycin and PMSF and imaged by widefield microscopy. Scale bar 1µm.

(B) Timelapse microscopy of IM expansion shows an increase in vesicle size with maintenance of a narrow rim. Cells expressing mNG-Atg8 were grown to logarithmic phase, treated for 1.5-2.5h with rapamycin, and imaged by widefield microscopy at indicated time points. Scale bar 1µm.

(C) Atg2 shows a spatially-constricted rim association pattern during IM expansion. Cells expressing sfTq2-Atg8 and Atg2-mNG were grown to logarithmic phase, treated for 1.5-3h with rapamycin, and imaged by widefield microscopy at indicated time-points. Scale bar 1µm.

#### Figure S2: The Atg2-Atg18 complex promotes constriction of the IM rim

(A) Auxin titration of Atg2-AID* leads to gradual loss of Atg2 expression and autophagic flux of GFP-Atg8. Cells of indicate genotype expressing GFP-Atg8 were grown to logarithmic phase, treated for 4h with rapamycin, and indicate the concentration of indole-3-acetic acid (IAA) and extracted proteins were immunoblotted as indicated. Error bars – S.E.M from 3 experiments, paired t-test: * p<0.05, *** p<0.0001, ns – non-significant.

(B) AID*-tagged Atg2 and Atg18 are depleted *in situ* from the IM rim. Cells of indicated genotype expressing sfTq2-Atg8 were grown to logarithmic phase, treated for 2-2.5h with rapamycin, PMSF, and indicated IAA concentration, and imaged by widefield microscopy. Scale bar 1µm.

(C) Core autophagy proteins are required for formation of IMs with open rims upon partial depletion of Atg2-AID*. Atg2-AID* cells of indicated genotype expressing mNG-Atg8, TagBFP-Pho8, mScarletI-PX$^{Vam7}$ were grown to logarithmic phase in presence of 10µM $CuSO_4$, treated for 2.5h with rapamycin, PMSF and 10µM IAA, and imaged by widefield microscopy. Scale bar 1µm.

(D) Canonical association pattern of autophagy proteins with IMs of open rim – Atg1 and Atg5 localize throughout the IM surface and at its base, respectively. Atg2-AID* cells expressing sfTq2-Atg8 and indicated mNG-tagged protein were grown to logarithmic phase, treated for 2h with rapamycin, PMSF, and 10µM IAA, and imaged by widefield microscopy. Scale bar 1µm.

(E) Correlative light electron microscopy (CLEM) analysis of partially-depleted Atg2 IMs shows double membranes with an open, locally-deformed Atg24-associated rim. Atg2-AID* cells expressing mNG-Atg8 and Atg24-mScarletI were grown to logarithmic phase and treated for 100min with rapamycin, PMSF, and 5µM IAA before CLEM processing and imaging. A single z-slice of reconstructed tomography z-stack is shown. Scale bar 200nm.

(F) Down-regulation of Atg2 by a weak version of the copper-induced promoter CUP1 leads to a severe loss of Atg2 expression levels and autophagic flux of GFP-Atg8 – mildly rescued by copper-induced expression. Cells of indicated genotype expressing GFP-Atg8 were grown to logarithmic phase in the presence of indicated $CuSO_4$ concentration, treated for 4h with rapamycin and extracted proteins were immunoblotted as indicated. Error bars – S.E.M from 3 experiments, paired t-test: ** $p<0.001$, *** $p<0.0001$, ns – non-significant.

#### Figure S3: An obligate complex of Atg24 with Atg20 or Snx41 assembles at the IM rim

(A, B) Atg20 and Snx41 C-terminally tagged with mScarletI localizes to the IM rim. Cells of indicated genotype expressing mNG-Atg8, TagBFP-Pho8, and Atg20-mScarletI (A) or Snx41-mScarletI (B) were grown to logarithmic phase, treated for 2.5-3.5h with rapamycin and PMSF, and imaged by widefield microscopy. Scale bar 1µm.

(C) Snx41 N-terminally tagged with mScarletI localizes to the IM rim. Cells of indicated genotype expressing mNG-Atg8, TagBFP-Pho8, and mScarletI-Snx41 were grown to logarithmic phase, treated for 2.5-4h with rapamycin and PMSF, as well as 250µM IAA for depletion of indicated AID*-tagged proteins, and imaged by widefield microscopy. Scale bar 1µm.

(D, E) Atg24 (D) and Atg20 (E) associate in a spatially-extended manner with the open IM rim upon partial depletion of Atg2-AID* in an Atg14/Atg18-dependent manner. Cells of indicated genotype expressing mNG-Atg8, TagBFP-Pho8, and mScarletI-Atg24 (D) or mScarletI-Atg20 (E) and were grown to logarithmic phase treated for 1.5-3h with rapamycin, PMSF and 10µM IAA and imaged by widefield microscopy. Scale bar 1µm.

(F) Atg24 associates stably with the open IM rim upon partial depletion of Atg2-AID*. Atg2-AID* cells expressing mNG-Atg8 and Atg24-mScarletI were grown to logarithmic phase, treated for 2-3h with rapamycin and 10µM IAA, and imaged by widefield microscopy at indicated time-points. Scale bar 1µm.

#### Figure S4: The Atg24 complex promotes spherical IM morphogenesis

(A) Core autophagy proteins are required for the formation of Atg24-deficient elongated IMs. Atg24-AID* cells of indicated genotype expressing mNG-Atg8, TagBFP-Pho8, mScarletI-PX$^{Vam7}$ were grown to logarithmic phase in presence of 10µM $CuSO_4$, treated for 2.5-3h with rapamycin, PMSF and 250µM IAA, and imaged by widefield microscopy. Scale bar 1µm.

(B) The morphogenic effect of Atg24 on IMs is observed in genetic backgrounds other than w303. Atg11-deficient cells of indicated BY4741 or SEY6210 background and genotype expressing mNG-Atg8 were grown to logarithmic phase and treated for 1.5-2h with rapamycin and PMSF before imaging by widefield microscopy. Scale bar 1µm.

(C) CLEM analysis of Atg24-deficient cells exhibits narrow tubular Atg8-positive IMs whose interior excludes cytosolic particles like ribosomes. Δatg24 Atg2-AID* cells expressing mNG-Atg8 were grown to logarithmic phase and treated for 100min with rapamycin, PMSF, and 5µM before CLEM processing and imaging. A single z-slice of reconstructed tomography z-stack is shown. Scale bar 200nm.

(D) Atg24 plays a quantitative morphogenic role in IM inflation. Δvps38Δymr1 Atg24-AID* cells expressing mNG-Atg8 and mScarletI-PX$^{Vam7}$ were grown to logarithmic phase, treated for 2-3h with rapamycin, PMSF, and indicated IAA concentration, and imaged by widefield microscopy. Scale bar 1µm.

#### Figure S5: Spatially-extended assembly of the Atg24 complex opens the IM rim

(A) The correlation between the spatial extension of Atg24 assembly and opening of the rim upon partial depletion of Atg2 is independent of endosomal PI3-kinase subunit Vps38 but depends on autophagic PI3-kinase subunit Atg14 and on rim recruitment of Atg24 by Atg20/Snx41. Atg2-AID* cells of indicated genotype expressing mNG-Atg8 and indicated Atg20 or Atg24 alleles were grown to logarithmic phase in presence of 10µM CuSO4 and treated with rapamycin, PMSF, and 10µM IAA for 2.5-4h, before imaging by widefield microscopy. Arrowheads point at cortical and nuclear ER or plasma membrane (PM) localization of Atg24 (see text). PM$^{Gpa2}$ – the PM-targeting acylation signal of Gpa2. TMD$^{Sec71}$ – the ER cytosol-facing targeting transmembrane domain of Sec71. ll – long linker. sl – short linker. Scale bar 1µm.

(B, C) The spatial extension of Atg24 or Atg20 assemblies correlates with rim opening even upon mock-treated, non-depleted Atg2-AID* conditions. Atg2-AID* cells of indicated genotype expressing mNG-Atg8 and indicated Atg24 (B) or Atg20 (C) alleles were grown to logarithmic phase in presence of 10µM $CuSO_4$ and treated with rapamycin, PMSF, and 0µM IAA for 2.5-3h, before imaging by widefield microscopy. ll – long linker. sl – short linker. Scale bar 1µm.

(D) The rim-opening activity of Atg20-sfCh3V is augmented compared with Atg20-mScarletI even in unperturbed *ATG2* wildtype conditions. Cells expressing mNG-Atg8 and indicated allele of Atg20 were grown to logarithmic phase, treated for 2-2.5h with rapamycin and PMSF, and imaged by widefield microscopy. Scale bar 1µm.

(E) The augmented rim-opening activity of Atg24-sfCh3V compared with Atg24-mScarletI is Atg2-dependent, Vps38-independent. Cells of indicated genotype expressing mNG-Atg8 and indicated allele of Atg24 were grown to logarithmic phase, treated for 2-2.5h with rapamycin and PMSF, and imaged by widefield microscopy. Scale bar 1µm.

(F, G) The augmented rim-opening activity of Atg24-sfCh3V compared with Atg24-mScarletI is observed in genetic backgrounds other than w303. Atg11-deficient cells of the BY4741 (F) or SEY6210 (G) background and indicated genotype expressing mNG-Atg8 and indicated Atg24 allele were grown to logarithmic phase and treated for 1.5-2h with rapamycin and PMSF before imaging by widefield microscopy. Scale bar 1µm.

#### Figure S6: Excessive activity of the Atg24 complex impairs IM maturation

(A, B) Partial depletion of Atg2-AID* leads to kinetic delay in autophagic body formation but largely maintains IM size. Atg2-AID* cells expressing mNG-Atg8 and TagBFP-Pho8 were grown to logarithmic phase in presence of 10µM $CuSO_4$, treated for indicated time with rapamycin, PMSF, and indicated IAA concentration, and imaged by widefield microscopy. Arrowheads point at autophagic bodies. Scale bar 1µm.

(C) Prolonged starvation coupled with partial depletion of Atg18-AID* leads to loss of vacuolar delivery of Atg8 and retention of a larger cytosolic IM – compared with wildtype. Cells of indicated genotype expressing mNG-Atg8, mScarletI-PX$^{Vam7,}$ and TagBFP-Pho8 were grown to logarithmic phase in presence of 10µM $CuSO_4$, washed into nitrogen starvation medium (SD-N), treated with 250µM IAA for 16h and imaged by widefield microscopy. Scale bar 1µm.

(D) The rim-augmenting activity of Atg24-sfCh3V leads to kinetic delay in autophagic body formation upon partial depletion of Atg2-AID* – in comparison with Atg24-mScarletI. Δatg15 Atg2-AID* cells expressing mNG-Atg8 and indicated Atg24 allele were grown to logarithmic phase, treated for indicated time with rapamycin, PMSF, and 10µM IAA concentration, and imaged by widefield microscopy. Arrowheads point at autophagic bodies. Scale bar 1µm.

(E) The rim-augmenting activity of Atg24-sfCh3V exacerbates the loss of GFP-Atg8 flux due to partial depletion of Atg2-AID* – compared with Atg24-mScarletI. Atg2-AID* cells expressing GFP-Atg8 and indicated Atg24 allele were grown to logarithmic phase, treated for 3.5h with rapamycin and extracted proteins were immunoblotted as indicated. Error bars – S.E.M from 3 experiments, paired t-test: * $p<0.05$, ** $p<0.001$, *** $p<0.0001$.

(F) The rim-augmenting Atg24-sfCh3V stably associates with the open IM rim upon partial depletion of Atg2-AID*. Atg2-AID* cells expressing mNG-Atg8 and Atg24-sfCh3V were grown to logarithmic phase, treated for 1.5-2.5h with rapamycin and 10µM IAA, and imaged by widefield microscopy at indicated time-points. Scale bar 1µm.

#### Figure S7: Atg24 couples IM maturation with sequestration of large cargo

(A) Atg24 is dispensable for Atg2-dependent, Vps38-independent formation of autophagic bodies including small cargo protein Fba1 – but essential for autophagy of large cargo proteins 60S ribosomal subunit Rpl9b and Fba1-decorated *Aquifex aeolicus* lumazine synthase (AaLS) nanoparticle. Δvps38 Atg24-AID* cells expressing indicated mScarletI-tagged cargo protein were grown to logarithmic phase, treated for 5-5.5h with rapamycin, PMSF, and mock (0µM) or saturating (250µM) IAA concentration for indicated basal or depleted Atg24 levels, respectively, and imaged by widefield microscopy. Arrowheads point at autophagic bodies. Scale bar 1µm.

(B) Tethering of large cargo proteins (40S subunit Rps1b and 60S subunit Rpl8a) to Atg8 does not rescue the loss of autophagic delivery to vacuolar autophagic bodies upon depletion of Atg24. Atg24-AID* cells expressing Atg8, tagged with the GFP derivative sfTq2 and indicated cargo protein, tagged with mNG fused to a GFP-binding protein (GBP), were grown to logarithmic phase. After treatment for 2-2.5h with rapamycin, PMSF, and mock (0µM) or saturating (250µM) IAA concentration for basal or depleted Atg24 levels, respectively, autophagic bodies and cargo proteins were visualized by widefield microscopy. Arrowheads point at autophagic bodies. Scale bar 1µm.

(C) Depletion of rim-localized Atg24-AID* leads to Ymr1-independent tubulation of the Atg2-dependent IM. Atg24-AID*-sfCh3V cells with indicated genotype expressing mNG-Atg8 and sfTq2-fused indicated cargo protein were grown to logarithmic phase, treated for 4-5h with rapamycin, PMSF, and mock (0µM) or saturating (250µM) IAA for indicated basal or depleted Atg24 levels, respectively, and imaged by widefield microscopy. Arrowheads point at autophagic bodies. Scale bar 1µm.

(D) Working model – rim aperture of autophagic membranes balances cargo inclusion with vesicle maturation. In wildtype conditions of basal Atg24 complex activity at the rim ("Basal"), nonselective IMs expand in the shape of amphoras with, as the vesicle body grows in a spherical manner while the rim is kept narrow. The Atg24 complex keeps the rim sufficiently wide for sequestration of large cargo and is cleared from the nonselective IM rim in due time by Ymr1-mediated turnover of PI(3) to facilitate maturation. Lost activity of the Atg24 complex at the nonselective IM rim ("Absent") leads to excessive constriction the rim. Consequently, sequestration of small cargo is maintained and the IMs elongate in a tubular shape, while sequestration of large cargo is excluded. Under these conditions maturation is independent of PI(3)P turnover by Ymr1. Excessive assembly of the Atg24 complex at the rim ("Augmented"), on the other hand, leads to expansion of an aberrantly-open nonselective IM, which may fit large cargo but fails to mature. Eventual clearance of rim-associated Atg24 by Ymr1-mediated turnover of PI(3)P may allow maturation. Color code: blue – IM, pink – small cargo, green – large cargo, orange – Atg24 complex (24C).

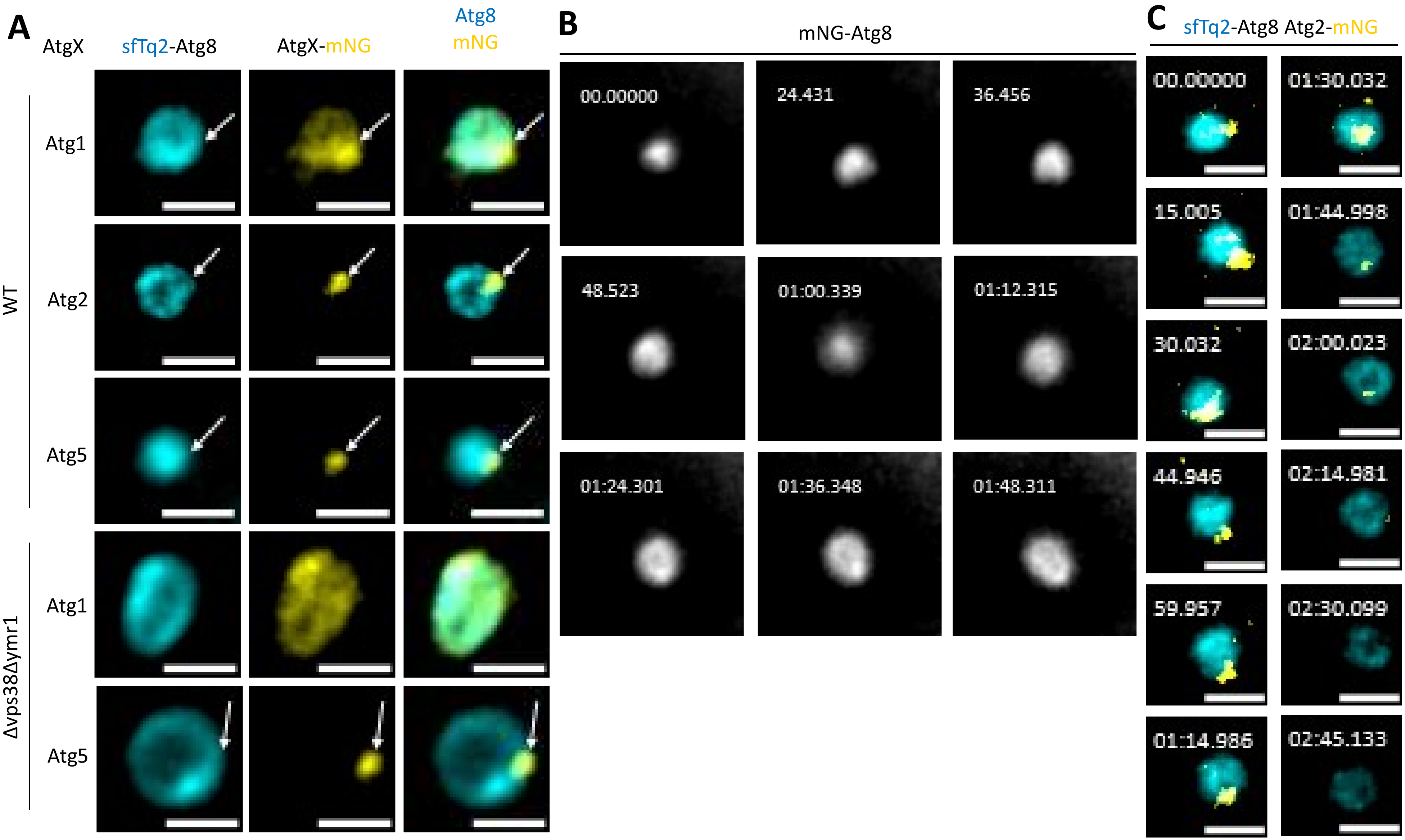
A
AtgX
sfTq2-Atg8
AtgX-mNG
Atg8
mNG
WT
Atg1
Atg2
Atg5
Δvps38Δymr1
Atg1
Atg5
B
mNG-Atg8
00.00000
24.431
36.456
48.523
01:00.339
01:12.315
01:24.301
01:36.348
01:48.311
C
sfTq2-Atg8 Atg2-mNG
00.00000
01:30.032
15.005
01:44.998
30.032
02:00.023
44.946
02:14.981
59.957
02:30.099
01:14.986
02:45.133

**Fig. S1**

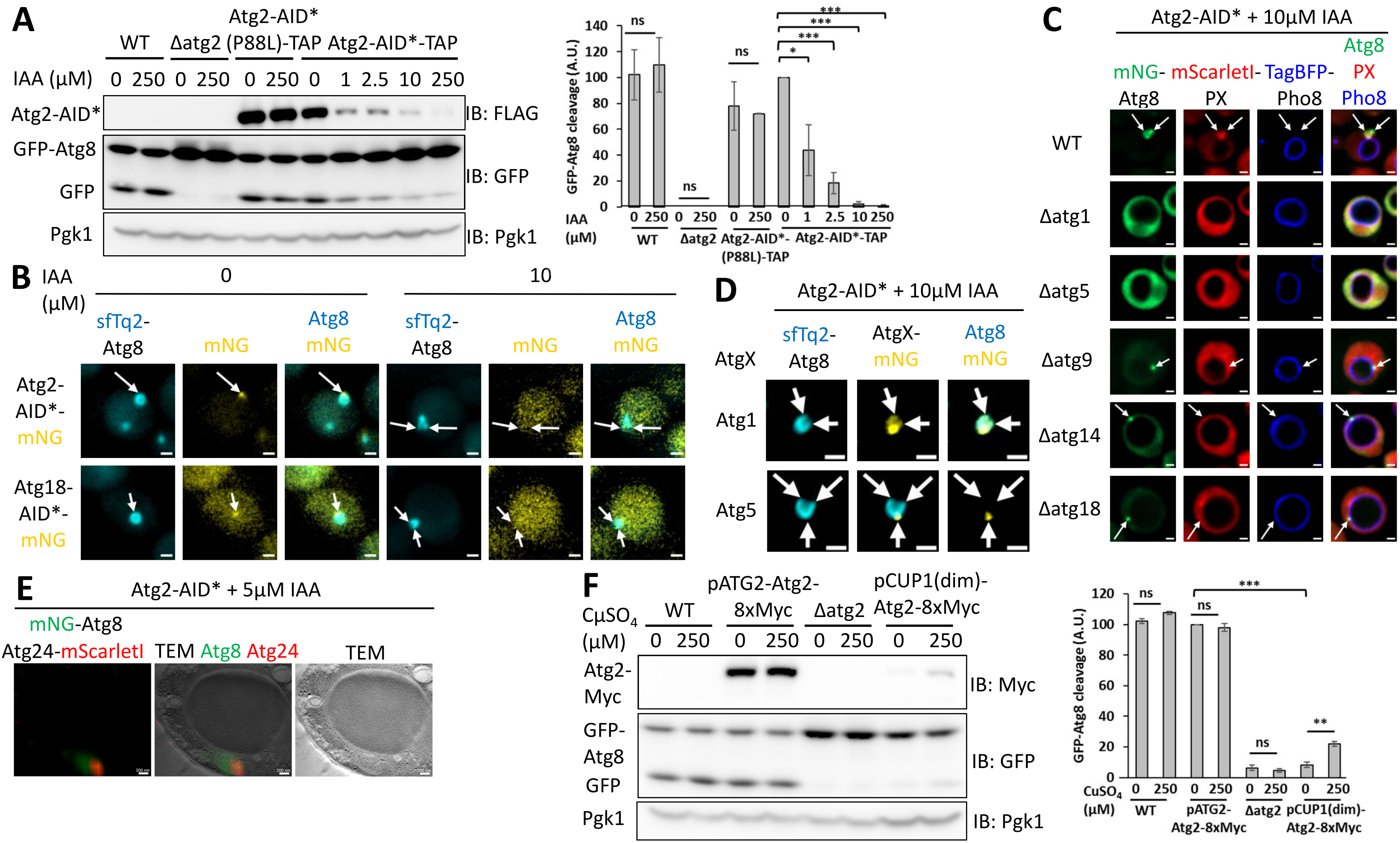
A
Atg2-AID*
WT
Δatg2
(P88L)-TAP
Atg2-AID*-TAP
IAA (μM)
0 250 0 250 0 250 0 1 2.5 10 250
Atg2-AID*
GFP-Atg8
GFP
Pgk1
IB: FLAG
IB: GFP
IB: Pgk1
GFP-Atg8 cleavage (A.U.)
ns
***
*
140
120
100
80
60
40
20
0
IAA (μM)
Atg2-AID*-(P88L)-TAP
B
IAA (μM)
0
10
sfTq2-Atg8
mNG
Atg8 mNG
Atg2-AID*-mNG
Atg18-AID*-mNG
C
Atg2-AID* + 10μM IAA
mNG-Atg8
mScarletI-PX
TagBFP-Pho8
Atg8 PX Pho8
WT
Δatg1
Δatg5
Δatg9
Δatg14
Δatg18
D
Atg2-AID* + 10μM IAA
AtgX
sfTq2-Atg8
AtgX-mNG
Atg8 mNG
Atg1
Atg5
E
Atg2-AID* + 5μM IAA
mNG-Atg8
Atg24-mScarletI
TEM Atg8 Atg24
TEM
F
CuSO4 (μM)
WT
pATG2-Atg2-8xMyc
Δatg2
pCUP1(dim)-Atg2-8xMyc
0 250 0 250 0 250 0 250
Atg2-Myc
GFP-Atg8
GFP
Pgk1
IB: Myc
IB: GFP
IB: Pgk1
GFP-Atg8 cleavage (A.U.)
ns
***
**
120
100
80
60
40
20
0
CuSO4 (μM)
pATG2-Atg2-8xMyc
pCUP1(dim)-Atg2-8xMyc

Fig. S2

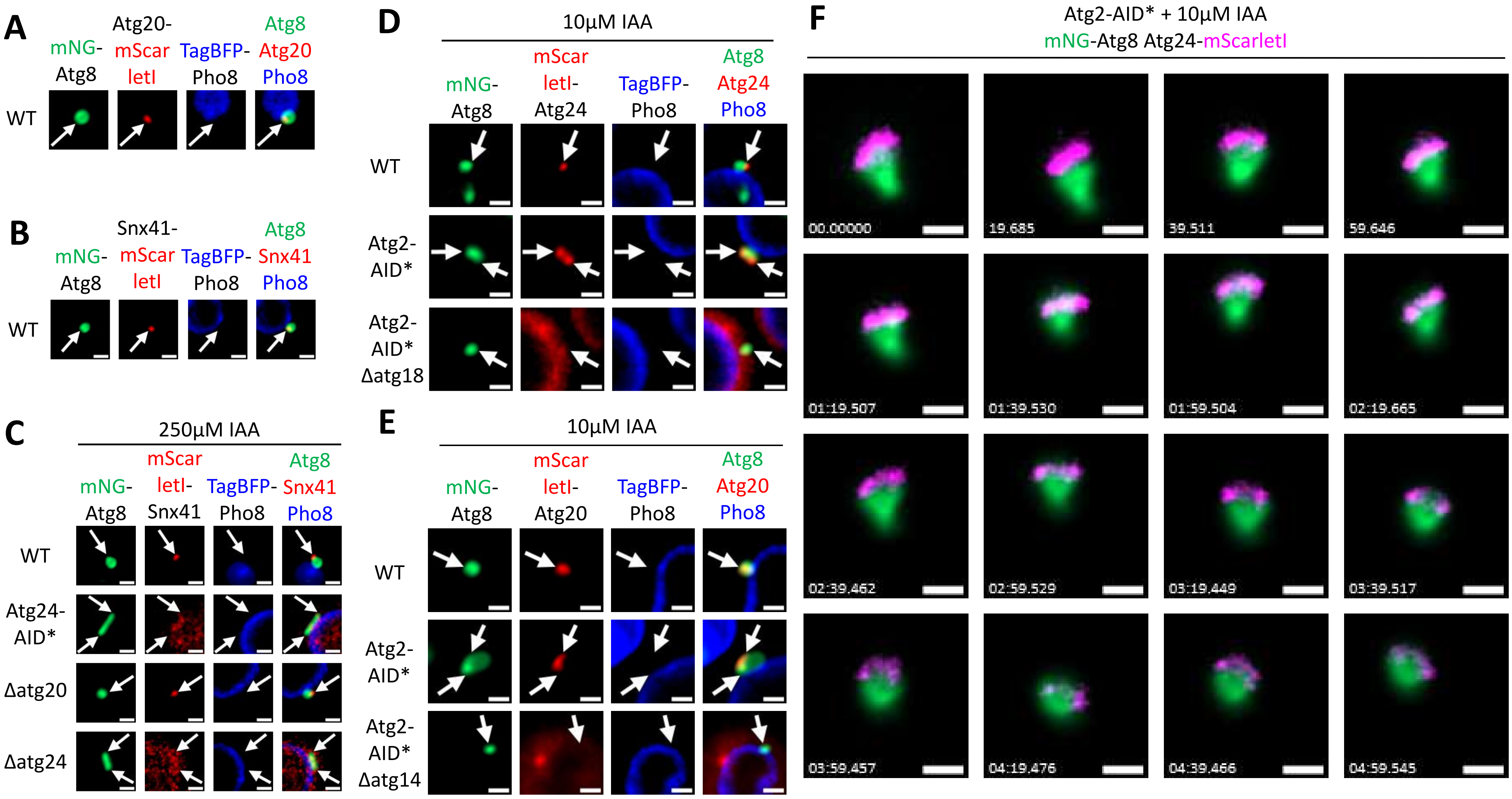
A
mNG-
Atg8
Atg20-
mScar
letI
TagBFP-
Pho8
Atg8
Atg20
Pho8
WT
B
mNG-
Atg8
Snx41-
mScar
letI
TagBFP-
Pho8
Atg8
Snx41
Pho8
WT
C
250μM IAA
mNG-
Atg8
mScar
letI-
Snx41
TagBFP-
Pho8
Atg8
Snx41
Pho8
WT
Atg24-
AID*
Δatg20
Δatg24
D
10μM IAA
mNG-
Atg8
mScar
letI-
Atg24
TagBFP-
Pho8
Atg8
Atg24
Pho8
WT
Atg2-
AID*
Atg2-
AID*
Δatg18
E
10μM IAA
mNG-
Atg8
mScar
letI-
Atg20
TagBFP-
Pho8
Atg8
Atg20
Pho8
WT
Atg2-
AID*
Atg2-
AID*
Δatg14
F
Atg2-AID* + 10μM IAA
mNG-Atg8 Atg24-mScarletI
00.00000
19.685
39.511
59.646
01:19.507
01:39.530
01:59.504
02:19.665
02:39.462
02:59.529
03:19.449
03:39.517
03:59.457
04:19.476
04:39.466
04:59.545

**Fig. S3**

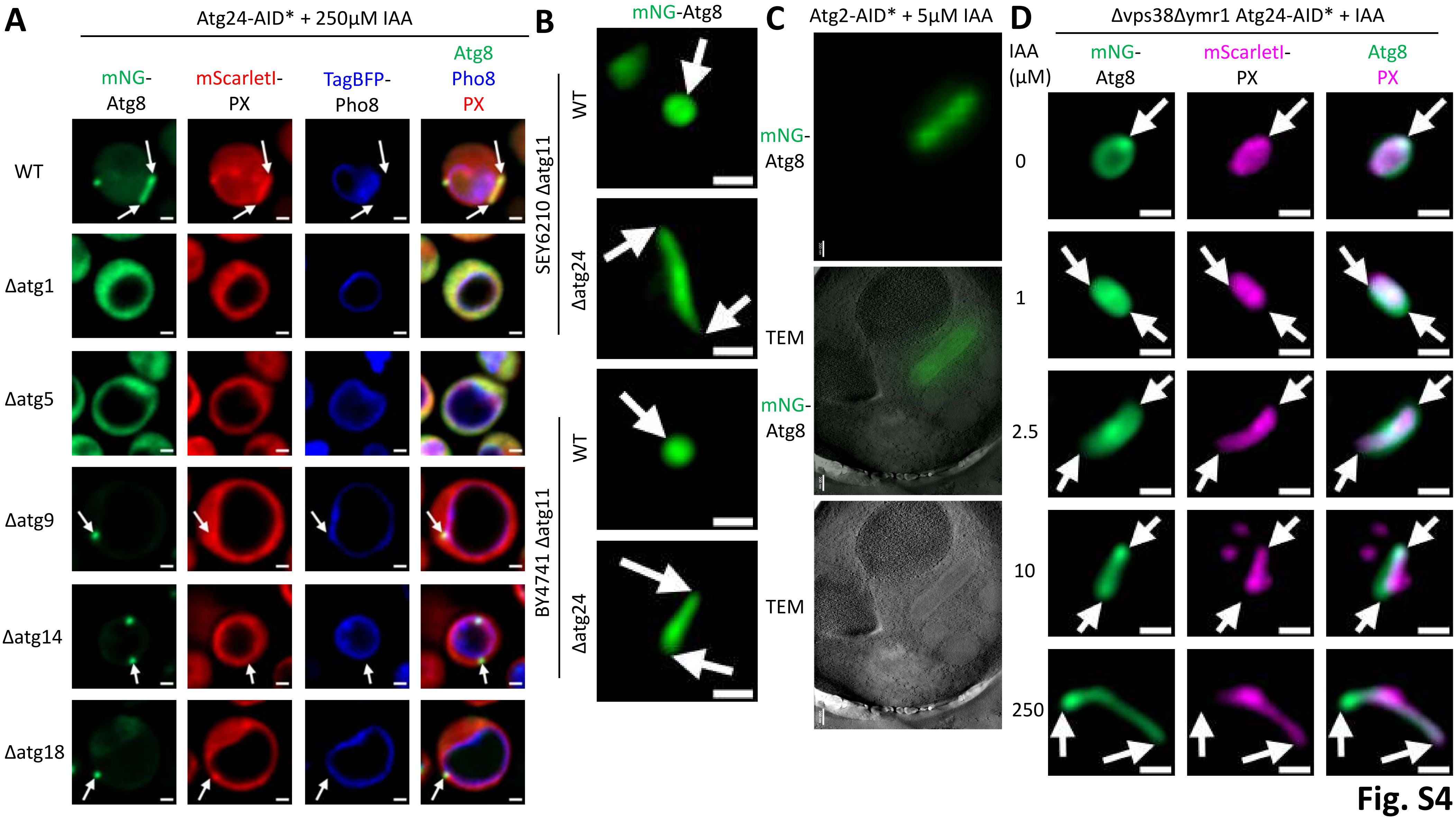
A
Atg24-AID* + 250µM IAA
mNG-Atg8
mScarletI-PX
TagBFP-Pho8
Atg8 Pho8 PX
WT
Δatg1
Δatg5
Δatg9
Δatg14
Δatg18
B
mNG-Atg8
SEY6210 Δatg11
WT
Δatg24
BY4741 Δatg11
WT
Δatg24
C
Atg2-AID* + 5µM IAA
mNG-Atg8
TEM
mNG-Atg8
TEM
D
Δvps38Δymr1 Atg24-AID* + IAA
IAA (µM)
mNG-Atg8
mScarletI-PX
Atg8 PX
0
1
2.5
10
250

**Fig. S4**

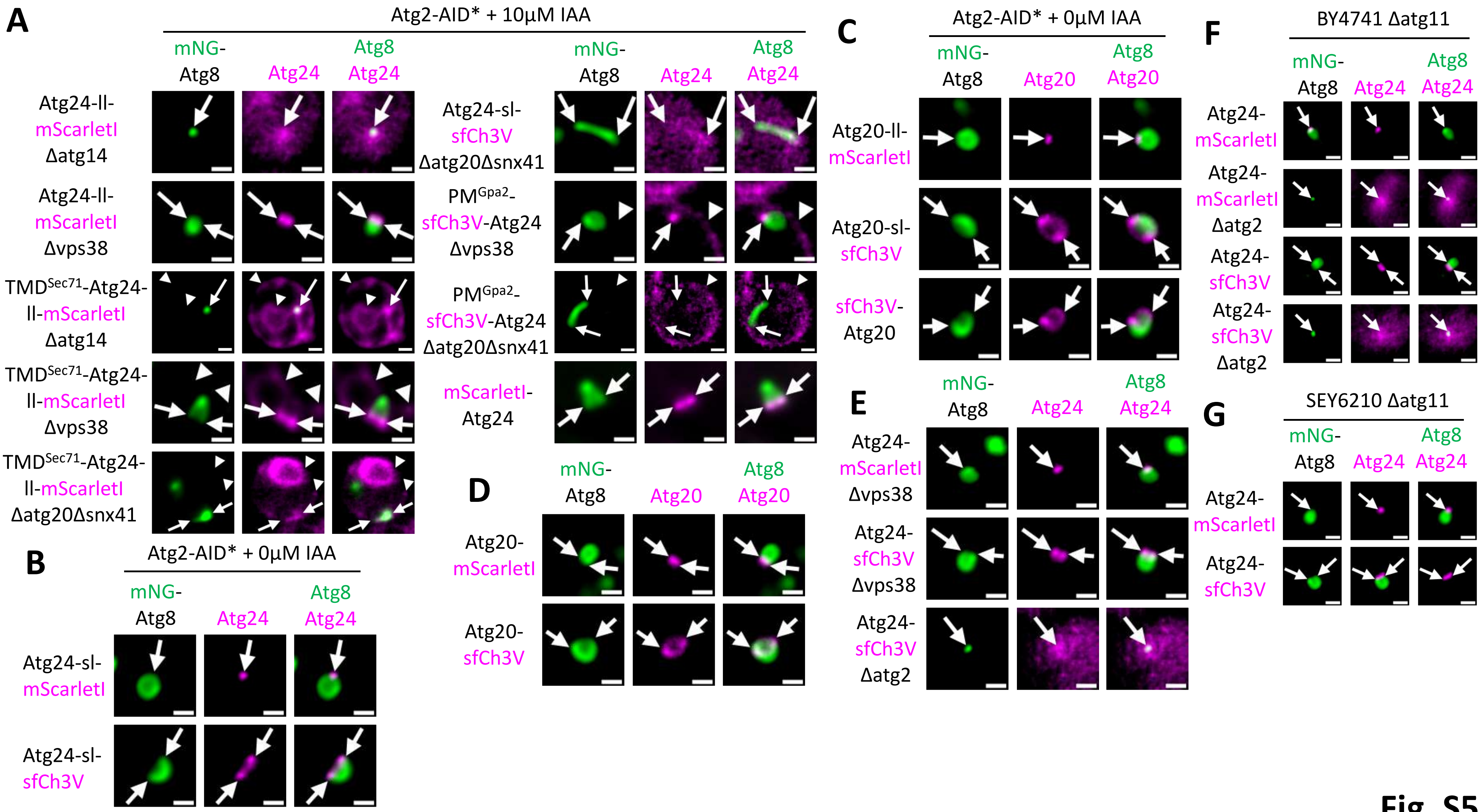
A
Atg2-AID* + 10µM IAA
mNG-Atg8
Atg24
Atg8 Atg24
Atg24-ll-mScarletI Δatg14
Atg24-ll-mScarletI Δvps38
TMDSec71-Atg24-ll-mScarletI Δatg14
TMDSec71-Atg24-ll-mScarletI Δvps38
TMDSec71-Atg24-ll-mScarletI Δatg20Δsnx41
Atg24-sl-sfCh3V Δatg20Δsnx41
PMGpa2-sfCh3V-Atg24 Δvps38
PMGpa2-sfCh3V-Atg24 Δatg20Δsnx41
mScarletI-Atg24
B
Atg2-AID* + 0µM IAA
Atg24-sl-mScarletI
Atg24-sl-sfCh3V
C
Atg2-AID* + 0µM IAA
Atg20
Atg8 Atg20
Atg20-ll-mScarletI
Atg20-sl-sfCh3V
sfCh3V-Atg20
D
Atg20-mScarletI
Atg20-sfCh3V
E
Atg24-mScarletI Δvps38
Atg24-sfCh3V Δvps38
Atg24-sfCh3V Δatg2
F
BY4741 Δatg11
Atg24-mScarletI
Atg24-mScarletI Δatg2
Atg24-sfCh3V
Atg24-sfCh3V Δatg2
G
SEY6210 Δatg11
Atg24-mScarletI
Atg24-sfCh3V

**Fig. S5**

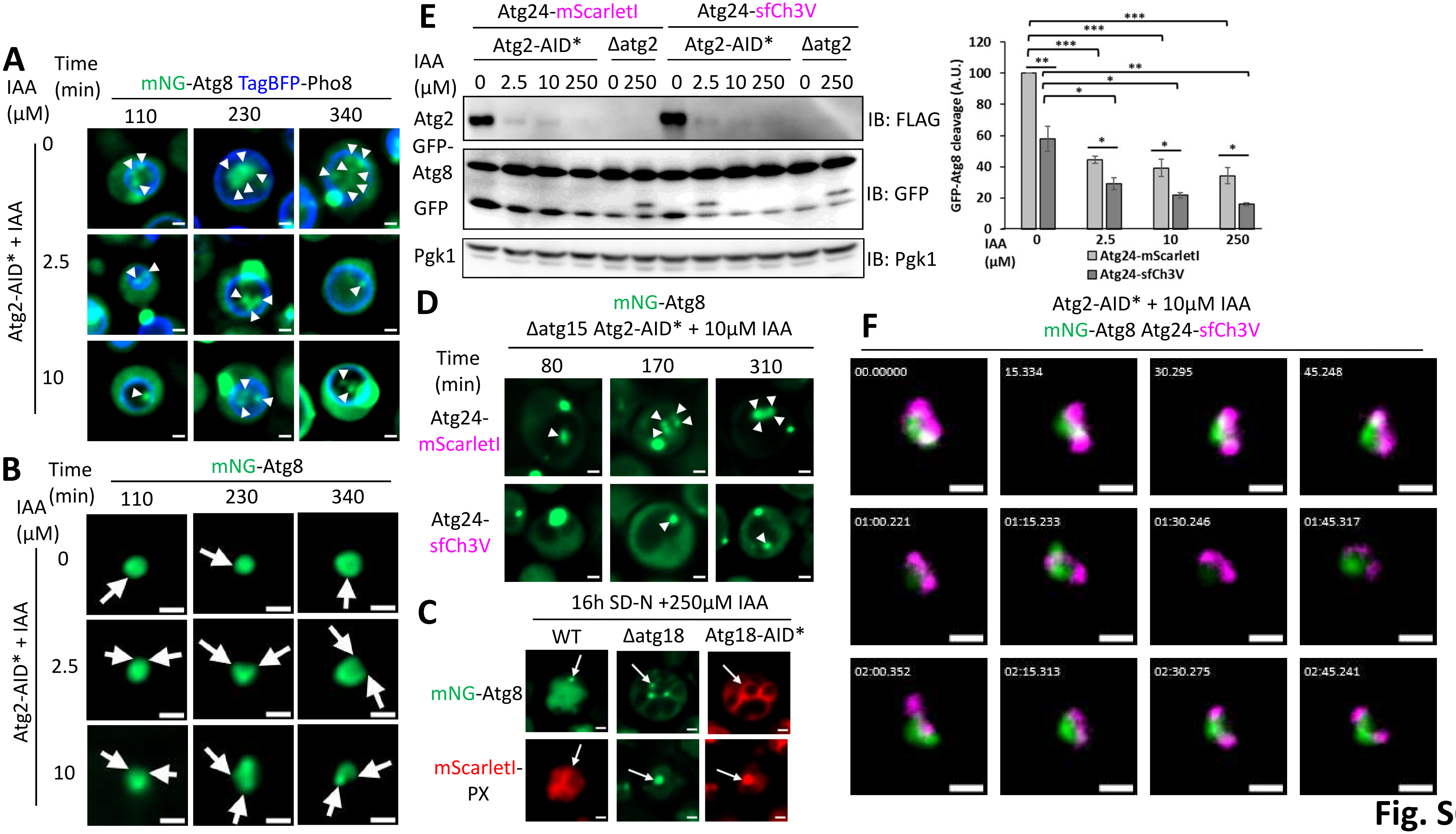
A
IAA (µM)
Time (min)
mNG-Atg8 TagBFP-Pho8
110
230
340
0
2.5
10
Atg2-AID* + IAA
B
Time (min)
IAA (µM)
mNG-Atg8
110
230
340
0
2.5
10
Atg2-AID* + IAA
E
Atg24-mScarletI
Atg24-sfCh3V
Atg2-AID*
Δatg2
Atg2-AID*
Δatg2
IAA (µM)
0 2.5 10 250 0 250 0 2.5 10 250 0 250
Atg2
GFP-Atg8
GFP
Pgk1
IB: FLAG
IB: GFP
IB: Pgk1
GFP-Atg8 cleavage (A.U.)
120
100
80
60
40
20
0
IAA (µM)
0
2.5
10
250
Atg24-mScarletI
Atg24-sfCh3V
***
***
***
**
**
*
*
*
*
*
*
D
mNG-Atg8
Δatg15 Atg2-AID* + 10µM IAA
Time (min)
80
170
310
Atg24-mScarletI
Atg24-sfCh3V
C
16h SD-N +250µM IAA
WT
Δatg18
Atg18-AID*
mNG-Atg8
mScarletI-PX
F
Atg2-AID* + 10µM IAA
mNG-Atg8 Atg24-sfCh3V
00.00000
15.334
30.295
45.248
01:00.221
01:15.233
01:30.246
01:45.317
02:00.352
02:15.313
02:30.275
02:45.241

**Fig. S6**

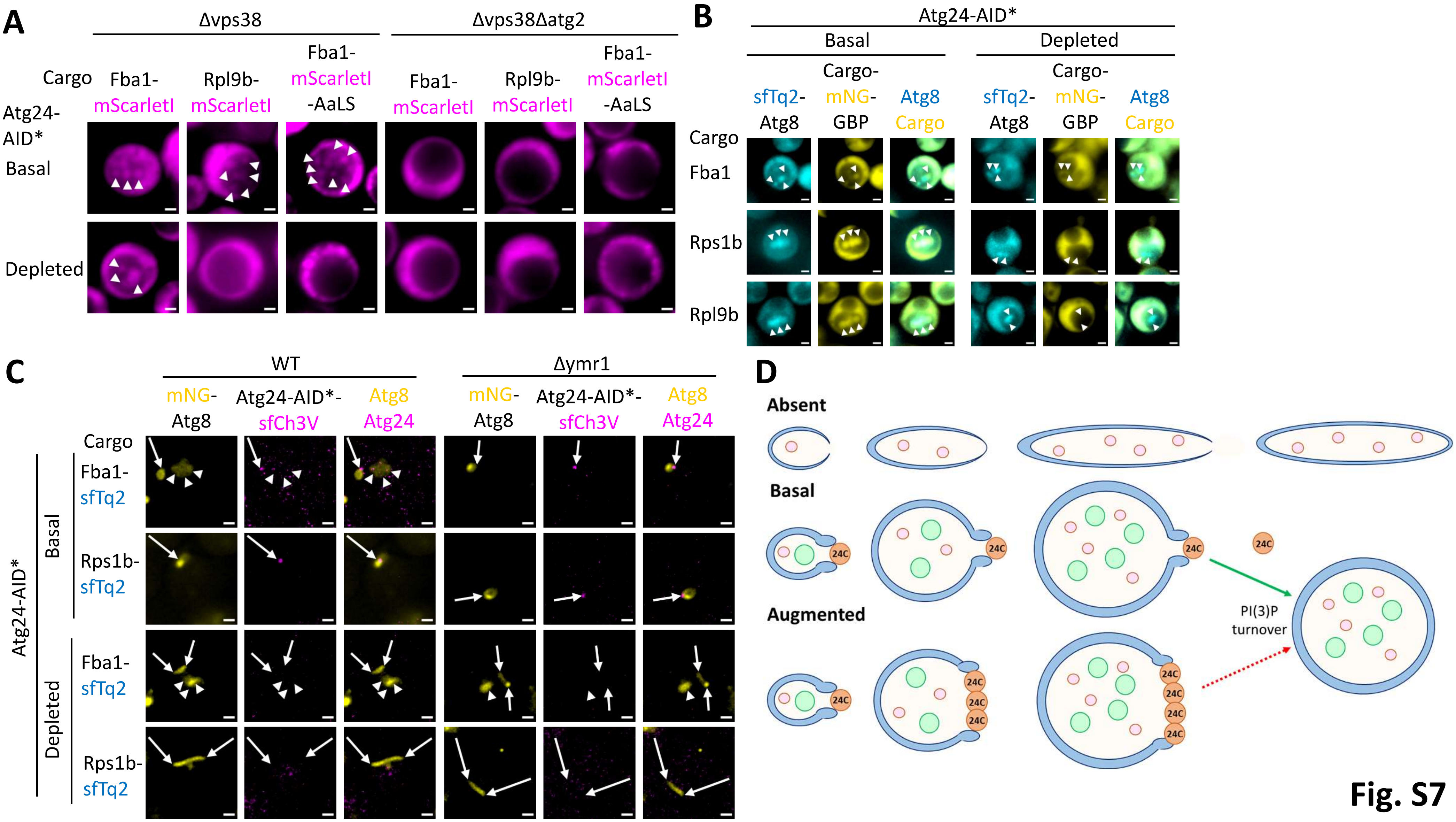
A
Δvps38
Δvps38Δatg2
Cargo
Fba1-mScarletI
Rpl9b-mScarletI
Fba1-mScarletI-AaLS
Atg24-AID*
Basal
Depleted
B
Atg24-AID*
Basal
Depleted
sfTq2-Atg8
Cargo-mNG-GBP
Atg8 Cargo
Cargo
Fba1
Rps1b
Rpl9b
C
WT
Δymr1
mNG-Atg8
Atg24-AID*-sfCh3V
Atg8 Atg24
Cargo
Atg24-AID*
Basal
Depleted
Fba1-sfTq2
Rps1b-sfTq2
D
Absent
Basal
Augmented
24C
PI(3)P turnover

Fig. S7
